# Phase Behavior and Dissociation Kinetics of Lamins in a Polymer Model of Progeria

**DOI:** 10.1101/2024.10.21.619497

**Authors:** Hadiya Abdul Hameed, Jaroslaw Paturej, Aykut Erbas

## Abstract

One of the key structural proteins in the eukaryotic cell nucleus is lamin. Lamins can assemble into a two-dimensional protein meshwork at the nuclear periphery, referred to as the nuclear lamina, which provides rigidity and shape to the nucleus. Mutations in lamin proteins that affect the structure of the nuclear lamina underlie laminopathic diseases, including Hutchinson–Gilford Progeria Syndrome (HGPS). Experiments have shown that, compared to healthy cells, lamin supramolecular structures (e.g., protofilaments) assemble into a thicker lamina structure in HGPS, where lamins form highly stable nematic microdomains at the nuclear periphery, reminiscent of liquid crystals. This significantly alters the morphological and mechanical properties of the nucleus. In this study, we investigate the aggregation of lamin fibrous structures and their dissociation kinetics from the nuclear periphery by modeling them as coarse-grained, rod-like polymer chains confined in a rigid spherical shell. Our model recapitulates the formation of multidirectional nematic domains at the nuclear surface and the reduced lamin dissociation observed in HGPS nuclei by adjusting lamin concentration, lamin-lamin (specifically head-tail), and lamin-shell association strengths. While nematic phase formation requires relatively strong lamin-shell affinity under any non-vanishing inter-lamin attraction, the thickness of this layer is primarily controlled by head-tail association strength in the model. Furthermore, the dissociation kinetics of lamin chains from the chain aggregates at the periphery (lamina) exhibits a concentration-dependent dissociation (facilitated dissociation) pattern governed by weak lamin-lamin interactions, reminiscent of healthy nuclei. Overall, our calculations demonstrate how an interplay between molecular interactions altered by mutations and lamin concentration can lead to an abnormal nuclear lamina in laminopathic diseases.

## I. INTRODUCTION

The nuclear lamina is a two-dimensional (2D) meshwork structure mainly composed of lamin proteins that are type V intermediate filaments [1–4]. The lamina is located at the nuclear periphery, underneath the inner nuclear membrane (INM)—the phospholipid bilayer of the nuclear envelope that faces the nucleoplasm—and maintains the shape and structural integrity of eukaryotic cell nuclei [5–10]. It is a highly dynamic meshwork, disassembling during the nuclear envelope breakdown stage of cell division into its molecular components and reassembling around the replicated decondensing chromosomes [11, 12]. This intricate process requires constituting lamin proteins to self-organize repeatedly and accurately at the nuclear periphery without interfering with the functional and mechanical properties of the nucleus.

The mammalian lamina comprises two main types of lamin proteins: A/C and B [2, 8]. Both lamin types share common structural features: a short N-terminal head domain, an *α*-helical central rod domain, and a globular tail domain [13–15]. The first level of structural assembly of lamin proteins is the formation of a dimer. This occurs when the central rod domains of two lamins coil in a parallel fashion [2, 14–16]. Dimers can further associate with one another to high-aspect ratio protofilaments or other higher-order fibrous structures [14, 15, 17] (see Supplementary Material Fig S1). Lamin fibers of various lengths and thicknesses sterically interact with each other and the INM [15, 18]. These reversible interactions allow lamin supramolecular structures to localize near the nuclear periphery, around the chromosomes, and form the nuclear lamina meshwork underneath the INM [13, 19, 20]. Both *in vitro* and *in vivo* studies suggest that the lamina is a relatively isotropic, heterogeneous network of lamin fibers in healthy cells, exhibiting polymer-network characteristics [21, 22] and “node” distributions [17, 22, 23]. Notably, in most healthy cells, lamin distribution at the nuclear lamina appears uniform at the scale of several microns under fluorescence microscopy [8, 22].

The molecular structure of the lamina is altered in a class of diseases collectively known as laminopathies. One such disease is Hutchinson–Gilford Progeria Syndrome (HGPS), or briefly Progeria, a rare genetic disease that causes the onset of premature aging [6, 8, 24–29]. The main cause of HGPS is a single-point mutation in the *LMNA* gene that encodes human lamin A [30–32]. This mutation produces a truncated form of lamin A, referred to as progerin, which lacks 50 amino acids at the tail domain. Progerin remains permanently farnesylated following post-translational modifications, unlike mature lamin A [24, 25, 27, 33–35]. Since farnesylation can enhance hydrophobic interactions, it can promote a stronger binding tendency of mutated lamin A (i.e., progerin) towards the INM or other hydrophobic components of the lamina. These alterations, in turn, could lead to anomalies in the morphology and mechanical properties of cell nuclei in Progeria, significantly decreasing cell viability [20, 36, 37].

In the progression of HGPS, as the nuclear concentration of mutant lamin A increases, the nucleus becomes more mechanically rigid and morphologically distorted as they accumulate at the nuclear lamina [34, 38–43]. Polarization light microscopy experiments further elucidated that lamins form orientally ordered, uncorrelated nematic domains in the nuclear lamina of HGPS nuclei, each large enough to display birefringence [7]. On the contrary, no birefringent pattern was observed at the lamina of healthy nuclei, suggesting the presence of randomly oriented lamin filaments [7, 41, 44]. Notably, studies also suggest that both wild-type lamin A (*wt* - lamin A) and progerin can form paracrystalline phases *in vitro* [15, 36], a property encountered in solutions of rod-like molecules with high bending rigidity (e.g., liquid crystals) [45–47]. More specifically, when confined by convex spherical surfaces, short (i.e., compared to the dimensions of the confining shell) polymer chains with high bending rigidity (e.g., rod-like) exhibit nematic phases in computational simulations [48–51]. This, combined with the available experimental data, indicates that the formation of nematic phase domains observed in nuclei of most laminopathic cells could result from irregular biomolecular interactions promoting increased localization of mutant lamins to the nuclear periphery.

Experiments further suggest that the dissociation properties of lamin proteins are altered in HGPS cells. Photobleaching experiments have shown that nucleoplasmic and peripheral lamins exchange with one another in healthy cells [7], further confirming that lamins can interact transiently with the lamina [52, 53]. In contrast to a healthy lamina, in Progeria, mutated lamin A, progerin, is irreversibly associated with the INM, exhibiting much weaker exchange kinetics with its solution counterparts compared to *wt* - lamin A [7, 26]. This indicates a potential correlation between lamina’s structural and kinetic properties in disease.

Overall, experimental evidence suggests that lamin proteins’ assembly and organization in the nuclear lamina are affected by lamin concentration and lamin affinity toward one another or the INM. Nevertheless, these factors’ relative biophysical roles in determining the nuclear lamina’s micro-scale properties, particularly in disease phenotypes, are relatively poorly understood. While a limited number of computational studies provide insights into the formation of misshapen nuclei, such as those observed in HGPS [9, 54], exploring physical interactions and stoichiometric conditions that can lead to the multi-directional nematic domains and thicker lamina in laminopathies can help decipher the universal principles of the disease progression.

While the nuclear lamina is a highly complex biomolecular structure, the formation of this protein meshwork in healthy cells requires: i) lamin localization near the INM [8]; ii) relatively stable association of lamins with one another to form polymer-network-like structures [22, 23]; iii) sufficient lamin concentration to occupy around 50 ± 10% of the inner nuclear surface [17]. To study these primary mechanisms involved in lamin assembly, we devise a simple molecular dynamics (MD) model for lamin fibers and consider them as a monodisperse solution of rod-like polymer chains confined in a spherical volume representing the nuclear void (Fig. 1). Focusing particularly on the effect of fiber concentration, head-tail association strength between fibers, and fiber-nuclear shell interactions, we emulate the formation of nematic phase domains and lamina thickening reported in HGPS while correlating slow lamin dissociation and lamina thickening observed in diseased nuclei. Our study suggests that the concentration alone cannot dictate lamin exchange kinetics or nuclear lamina thickening. Instead, there also needs to be a compromise between the strength of the inter-lamin association and the interaction of lamin fibers with the INM.

**FIG. 1.**
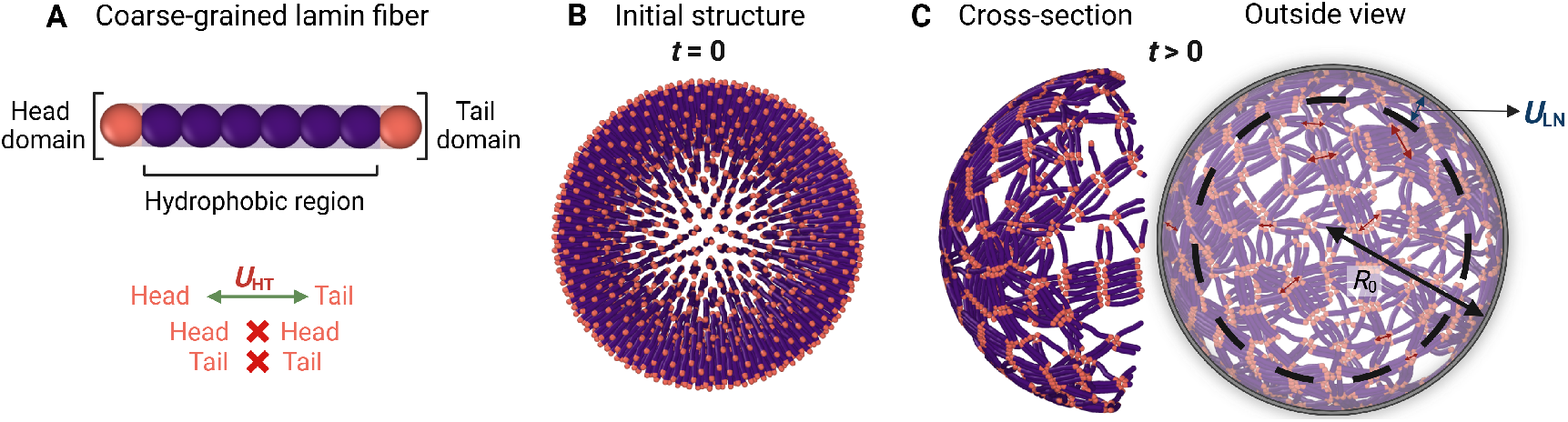
The schematics of the MD simulation model and simulation steps. (*A*) Coarse-grained lamin fiber with terminal head, tail, and central hydrophobic region. The terminal (head and tail) domains are allowed to associate with one another and are marked with the same color for simplification. (*B-C*) Illustration of the MD simulation steps. (*B*) The initial structure is minimized for 10^4^ MD time steps. (*C*) After minimization, the head-tail association potential, *U*_HT_, and lamin-shell interaction, *U*_LN_, are switched on and the simulation runs for another 10^6^ MD time steps.

## II. METHOD

### A. Molecular model of nuclear lamins

In the nucleoplasm, individual lamin proteins are not solvated but rather exist in their dimer form [2, 8]. On the contrary, lamins dimers typically aggregate near the nuclear boundary as more complex fibrous structures (e.g., short protofilaments) with high-aspect ratios, i.e, a lamina thickness of 14 ± 2 nm versus a mean fiber length of 380 ± 122 nm [17, 43]. Thus, in our study, each lamin fiber is approximated as a Kremer-Grest (KG) rod-like, bead-spring chain in implicit solvent (Fig. 1*A*) [55].

Given that HGPS and other laminopathies are predominantly associated with mutations in A-type lamins rather than B-type, we focus on modeling a single type of lamin fiber [30]. A-type lamins also play a key role in regulating the nuclear shape and stiffness [28, 31, 56]. Each lamin chain consists of head and tail domains, representing the terminal ends of lamin supramolecular structures (e.g., protofilaments). The central beads are repulsive, representing hydrophobic interactions at the core of lamin fibers (Fig. 1*A*). The beads representing the head, tail, and central hydrophobic region are all the same size, with each lamin fiber consisting of *n* = 8 beads unless otherwise specified. Using shorter or longer chains (i.e., *n* = 4 or *n* = 16) does not affect the results qualitatively (see Supplementary Material Fig. S2-S7). The lamin fibers are confined within a rigid spherical boundary of radius, *R*_0_, to mimic the steric effect of the nuclear envelope (Fig. 1*C*). The spherical confinement has a radius, *R*_0_ = 34*σ*, unless specified otherwise (Fig. 1). Here, the bead size is set to *b* = 1*σ*, which defines the unit length scale in simulations.

The concentration of lamin proteins (monomeric) was reported to be around *>* 100 nM *in vivo* [57]. Hence, it is difficult to estimate the quantity of lamin fibers (composed of many monomers) accumulating near the periphery. Thus, we calculated a polymer physical quantity, a 2-dimensional analog of concentration for fibers, by assuming a nuclear diameter of *d* ≈ 10 *μ*m and an approximate lamin fiber length of *l* ≈ 400 nm [5, 56, 58] to estimate the number of lamins required to cover the surface of as *N* ≈ (10 *μm/*4 × 10^2^ nm)^2^ ≈ 10^3^. To probe various concentration regimes, we varied lamin fiber concentration, *c*, by increasing the number of rods within our rigid, spherical nucleus model between *N* = 500 and *N* = 4000, covering between 50% to over 100% of the surface area of the confinement to account for the accumulation of progerin in HGPS nuclei and nucleoplasmic lamins in healthy cells [8, 17, 26]. This range also corresponds to lamin concentrations above and below the bulk overlap concentration, *c*\* (Eqn. 1). The bulk overlap concentration, *c*\*, represents the transition of a polymer solution from a dilute to a semi-dilute regime. This concentration marks where individual lamin chains begin to overlap and interact with one another. Notably, *c* = 1.6*c*\* i.e. *c* ≈ *c*\*, also corresponds to an occupied lamina volume of 13.5% in our simulations (Fig. 4) [17].

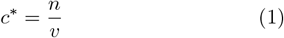

where *v* ≈ *l*^3^ *n*^3^ ≈ *σ*^3^ is the pervaded volume of a single lamin rod. The overlap concentration for 8-bead lamin rods, *n* = 8, is *c*\* ≈ 0.03*σ*^−3^. The concentration values used in the simulations are shown in Table I. We discuss our results by either using absolute concentration, *c*, with the units *σ*^−3^, or concentration normalized by *c*\*.

**Table I.** Conversion of number of chains (*N*) to concentration (*c* in *σ*^−3^ and *c* in terms of *c*\*)

| $N$ | $c(\sigma^{-3})$ | $c/c^*$ |
| --- | --- | --- |
| 500 | 0.02 | 0.8 |
| 750 | 0.04 | 1.2 |
| 1000 | 0.05 | 1.6 |
| 1250 | 0.06 | 2.0 |
| 1500 | 0.07 | 2.4 |
| 2000 | 0.097 | 3.3 |
| 2500 | 0.12 | 4.0 |
| 3000 | 0.15 | 4.8 |
| 3500 | 0.17 | 5.6 |
| 4000 | 0.19 | 6.5 |

### B. Simulation setup

In simulations, lamin rods are initially placed randomly within the spherical confinement (Fig. 1*B*). We verified that the initial configuration does not affect simulation results (see Supplementary Material Fig. S8). We minimize the energy of the initial lamin configuration for 10^4^ MD time steps (Fig. 1*B*) using an integration time step of Δ*t* = 0.001*τ*, where *τ* is the unit of time in the simulations. Following the minimization, each system undergoes an additional 10^6^ MD time steps with Δ*t* = 0.005*τ*, allowing the system to reach equilibrium (see Supplementary Material Fig. S9). During the production phase of the simulations, attractive lamin-shell and lamin-lamin interactions are introduced to the relaxed configuration to replicate healthy and diseased nuclear lamina phenotypes (Fig. 1*C*). The spherical confinement exerts an attractive potential, *U*_LN_, mimicking the interaction between the lamin fibers and the nuclear periphery (e.g., with the INM) (Fig. 1*C*). Since lamin proteins can assemble via head-tail interactions [2, 15], the head domain of each lamin fiber interacts exclusively with the tail domain of other lamins through a head-tail association potential, *U*_HT_ (Fig. 1*A*).

The non-bonded, steric interactions between chain beads and/ or with the inner confining surface are modeled by using the Lennard-Jones (LJ) potential:

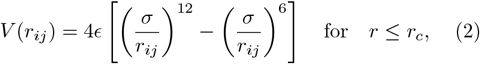

where *ϵ* represents the energy unit, *r*_*ij*_ is the distance between two beads, and *r*_*c*_ is the cut-off distance beyond which the LJ potential becomes negligible. For attractive interactions, such as head-tail and lamin-shell interactions, *r*_*c*_ is set to 2.5*σ*. For all other interactions, the cut-off radius is set to *r*_*c*_ = 2^1*/*6^*σ* to ensure steric repulsion. The default value of the interaction strength in Eq. 2 is *ϵ* = 1*k*_B_*T*, where *k*_B_ is the Boltzmann constant and *T* is the reduced absolute temperature, i.e., unity in our simulations. The attraction between the head and tail beads of lamin chains, *U*_HT_, as well as between lamins and the confining, structureless, surface, *U*_LN_, is adjusted by increasing *ϵ* in Eq. 2. Since affinities for lamin structures are not known, to probe various affinity ranges for *U*_HT_ and *U*_LN_, we varied attraction strengths as low as several *k*_B_*T* (susceptible to thermal fluctuations with energies on the order of *k*_B_*T*), and as high as *U*_LN_ = 10.0*k*_B_*T*. These attraction ranges are consistent with previous models with similar coarse-graining levels [59, 60].

All bonded interactions between adjacent beads in each lamin fiber are described using a Finitely Extensible Non-linear Elastic (FENE) potential:

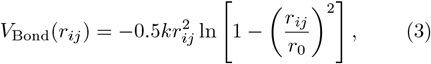

where the bond energy is set to *k* = 30.0*k*_B_*T/σ*^2^ and the maximum bond distance is set to *r*_0_ = 1.5*σ*.

The bending fluctuations of lamin chains are constrained by a harmonic angle potential in the form of:

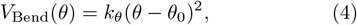

where the bending energy is set to *k*_*θ*_ = 10.0*k*_B_*T/rad*^2^, and the reference angle is *θ*_0_ = *π*. These parameters result in a persistence length higher than the length of each lamin chain, *l*_*p*_ *> l*, ensuring that they behave predominantly as semi-flexible polymers in the simulations (see Supplementary Material Fig. S10). All energies are defined in thermal energy units, *k*_B_*T* (≈ 0.6 *kcal/mol*).

All simulations were carried out in the LAMMPS MD simulations package [61]. Various Python libraries are used for data analysis [62]. VMD and OVITO were used for structural visualization [63, 64].

### C. Kinetic calculations

To quantify the kinetic behavior of lamin rods, we monitor the dissociation of peripherally localized chains across various binding affinities (*U*_HT_ and *U*_LN_) and lamin concentrations. Specifically, we measure the decay of the population of peripherally *bound* lamin fibers, while those freely diffusing in the nucleoplasm are considered *unbound*. As the simulation progresses, the initially bound lamins, *n*_0_, dissociate and exchange with those in the nucleoplasm. The number of remaining bound lamins, *n*(*t*), is calculated as a function of the time, *t* (see Supplementary Material Fig. S11). A lamin is considered dissociated and tagged as unbound if it moves beyond a defined cut-off radius, *R*_C_, from the nuclear periphery. If a dissociated lamin protein rebinds to the spherical confinement, it is not counted again as part of the surviving lamins, *n*(*t*). The dissociation cut-off distance is set to the size of a single chain, *R*_0_ *< R*_C_ ≤ 8*σ* [60, 65] (see Supplementary Material Fig. S12), and varies with the thickness, *δ*, of the nuclear lamina, which depends on lamin concentration. We also calculate the time-averaged concentration of *unbound*, free lamins, *c*_free_, as the average number of lamin rods that remain unbound over simulation time (see Supplementary Material). For the analysis, only the latter half of each simulation is considered, excluding the initial equilibration phase of lamin organization. A single-exponential fit is applied to model lamin dissociation and determine off-rates:

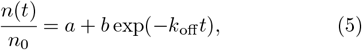

where *k*_off_ is the off-rate with the units of inverse simulation time, while *a* and *b* are fitting parameters. The fit is specifically applied when the bound lamin fraction satisfies *n*(*t*)*/n*_0_ ≤ 0.5, ensuring that it captures the later stages of dissociation, where the exponential decay behavior is most prominent.

## III. RESULTS

In our simulations, most lamin rods first localize near the spherical boundary at *t >* 0, forming a layer of adsorbed chains (Fig. 1*C*). We will refer to this layer as “lamina”. Once some or all of the chains are localized to the periphery, this is followed by either lamin dissociation or stable accumulation of chains to the spherical boundary depending on the lamin concentration, *c*, head-to-tail association strength, *U*_HT_, and the interaction of lamin rods with the interior surface of our spherical confinement, *U*_LN_.

### A. Nematic phase formation of model lamin chains on the spherical surface occurs beyond the threshold concentration, *c*\*

To explore the molecular conditions affecting the morphology and organization of lamin proteins in nuclear lamina assembly, we first analyze the effect of lamin chain concentration in a model nucleus in our simulations. We scan the concentrations above and below the threshold concentration, *c*\*, (Eq. 1), where lamin chains can be at steric contact in the bulk, 0.8*c*\* ≤ *c* ≤ 3.3*c*\* (Table I). Since we compare two interactions in our model, *U*_LN_ and *U*_HT_, we investigate their relative contribution to the nuclear lamina assembly process systematically. To do so, we first choose a high lamin-shell interaction strength, *U*_LN_ = 10.0*k*_B_*T*, while changing the head-tail association potential (Fig. 2). This ensures the localization of lamin chains near the boundary of the spherical confinement while allowing us to interrogate the effects of specific attractions between lamin fibers. For the head-tail attraction strength, values weaker than and equal to the lamin-shell attraction strength but higher than the thermal energy scale (i.e., ≈ *k*_B_ *T*) were chosen.

**FIG. 2.**
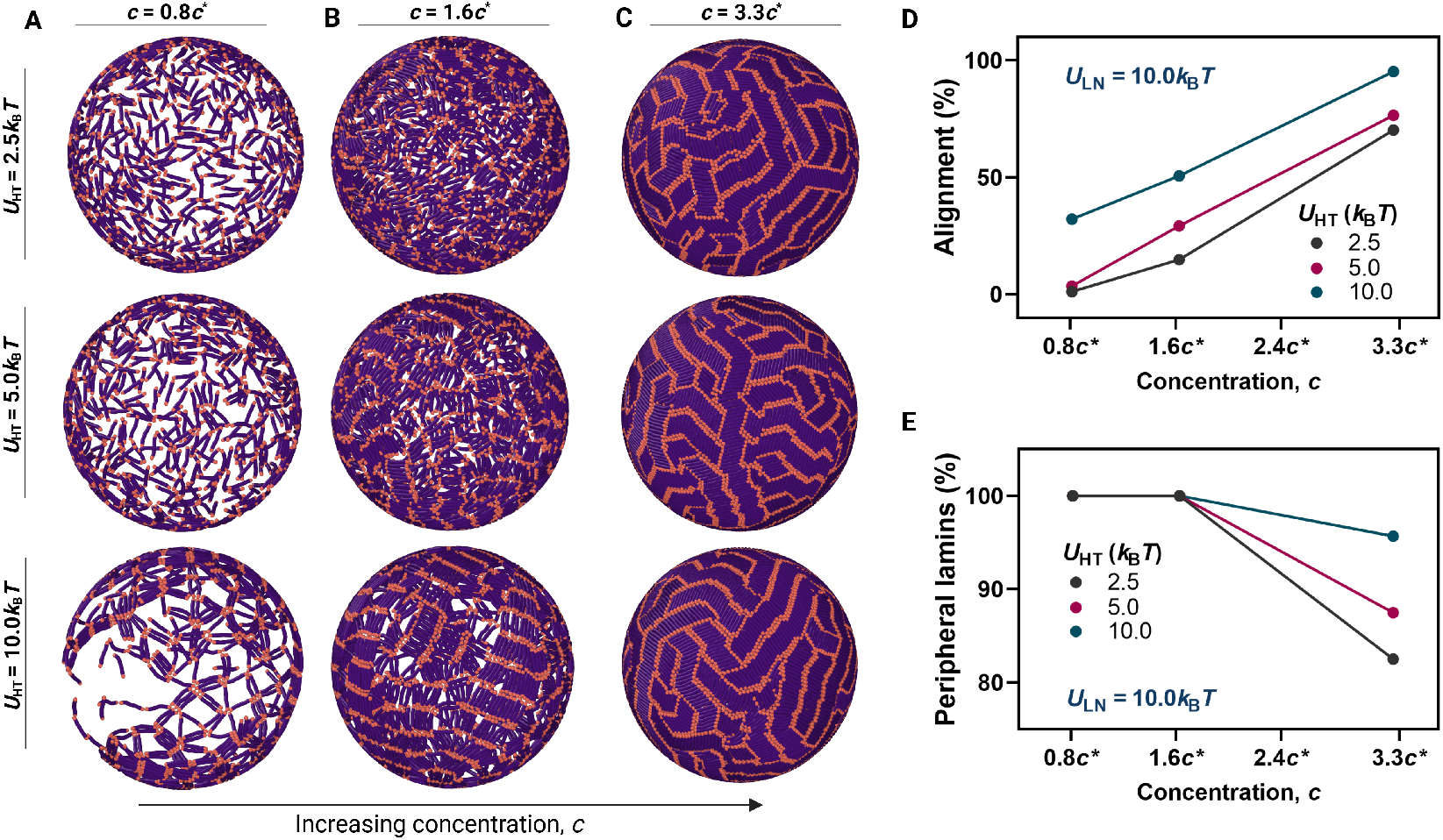
Isotropic and nematic phase formation of lamin chains at the nuclear lamina surface. (*A–C*) Representative snapshots of the rigid, model nucleus at various head-to-tail association potentials, *U*_HT_, with lamin concentrations, *c* = (*A*) 0.8*c*\*, (*B*) 1.6*c*\*, and (*C*) 3.3*c*\* at a fixed lamin-shell interaction strength, *U*_LN_ = 10.0*k*_B_*T*. (*D*) Percentage of lamin rod pairs aligned parallel to each other as a function of concentration, *c*, for various head-tail association strengths, *U*_HT_. (*E*) Percentage of peripheral lamin chains within a distance of 2.5*σ* from the inner surface for various head-tail association potentials, *U*_HT_.

Fig. 2 shows representative simulation snapshots taken from the final stage of different simulations at various concentrations. These simulations reveal that concentration directly affects the morphology of lamin phases localized at the inner surface of the model nucleus, particularly when lamin chains have a strong affinity towards the confining boundary, *U*_LN_ = 10.0*k*_B_*T*. At low concentrations (i.e., *c* ≤ *c*\*), lamin chains are arranged with no specific directional order regardless of the head-tail attraction strength (Fig. 2*A*, top to bottom). Notably, increasing the head-tail association strength does not affect their isotropic orientation, except the formation of nodes leading to a network configuration at *U*_HT_ = 10.0*k*_B_*T* (Fig. 2*A*, bottom). At intermediate to high concentrations, *c > c*\*, nematic-phase domains appear on the surface and cover the entire inner surface of the spherical confinement (Fig. 2*B*). These phases become more distinct as inter-lamin interaction is increased to *c >> c*\*, independent of *U*_HT_ (Fig. 2*C*). While these phases differ from global phases of semi-flexible chains reported earlier [49–51] and are visually non-percolating, they are consistent with the nematically ordered domains reported *in vivo* and *in vitro* for Progeria nuclei [6, 7, 36].

To further understand this nematic phase formation on the surface quantitatively, we attempt to use methodologies previously used to study nematic liquid crystalline phases of semi-flexible polymers [48, 66]. Accordingly, the diagonalization of the configurational tensor for all the chains shows whether they form a global nematic order in bulk or on the surface [48–51]. However, in our system, where the surface is covered by randomly oriented nematic domains (Fig. 2), this method cannot distinguish between nematic order and isotropic regions. Thus, we calculate a spin-spin-like order parameter instead, such that if a chain aligns next to its nearest neighbor, the order parameter is *S* = 1 or; otherwise *S* = 0. Averaging over pairs obeying the nearest neighbor criterium (*r*_nm_ *<* 1.5*σ*) as percentage alignment on the surface demonstrates the dependence of lamin ordering on the concentration and head-tail affinity (Fig. 2*D*). We also calculate the percentage of lamin chains within 2.5*σ* of the model nuclear confinement to quantify the effect of lamin-lamin interaction on nematic phase formation (Fig. 2*E*). Our results suggest that as both concentration, *c*, and head-tail interaction, *U*_HT_, increase, lamin alignment increases (Fig. 2*D*), as seen visually in Fig. 2*C*. This observation also seems to correlate to an increase in the peripheral lamins (Fig. 2*E*). We observe similar alignment trends for shorter lamin fibers (see Supplementary material Fig. S13, S14). These analyses suggest that while concentrations significantly above *c*\* lead to nematic domains, once chains are localized to the surface, the head-tail association strength controls the extent of this domain formation. In fact, at *U*_HT_ = 0, no phases are observed on the surface (see Supplementary material Fig. S15).

Overall, these simulations suggest that the strong localization of lamins on the nuclear surface can exhibit a transition from isotropic to nematic domains with increasing concentration of lamin species.

### B. Nematic phase formation on the shell surface is a result of the competition between lamin-lamin and lamin-shell interactions

Having demonstrated nematic phase formation via strong peripheral localization of lamin chains (i.e., *U*_LN_ ≥ *U*_HT_) (Fig. 2), we next explore whether a weaker *U*_LN_ can affect phase formation. In this investigation, a concentration slightly above our intermediate concentration (i.e., *c* = 2.0*c*\*) is chosen to ensure that we observe nematic phases (Fig. 2 and see Supplementary material Fig. S16).

In Fig. 3, from left to right, we increase the lamin-lamin (head-tail) attraction, *U*_HT_, while lamin-shell attraction, *U*_LN_, is increased from top to bottom in separate simulations.

**FIG. 3.**
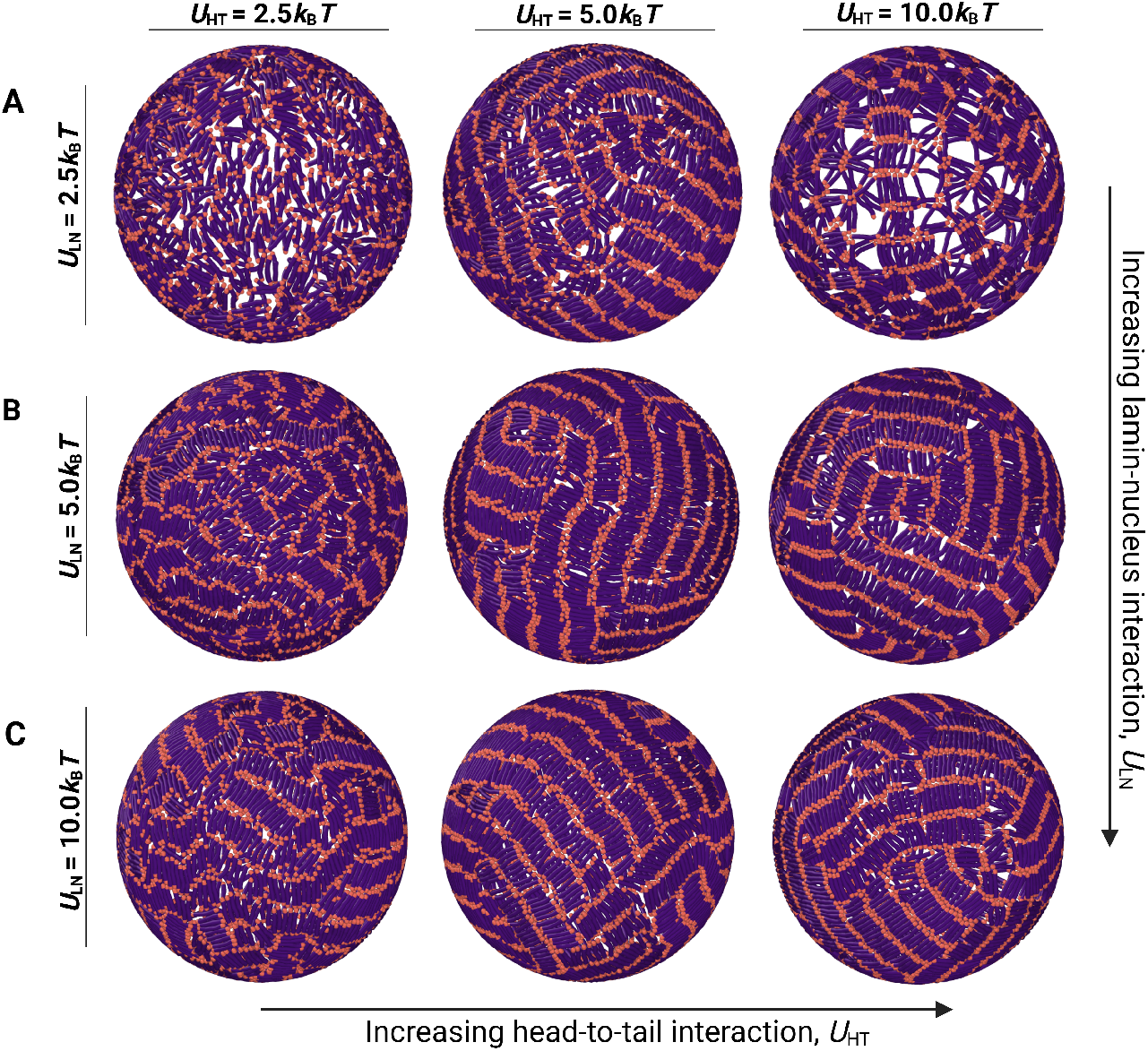
Representative snapshots of the exterior of the shell for various values of *U*_LN_ and *U*_HT_ at a concentration, *c* = 2.0*c*\*. (*A-C*) From top to bottom, *U*_LN_ increases. *U*_HT_ increases from left to right. Only those chains within a distance of 2.5*σ* from the surface are shown, while those in the interior of the confinement are not shown for clarity.

The weakest lamin-shell association strength used (i.e., *U*_LN_ = 2.5*k*_B_*T*) leads to either isotropic chain organization or a disrupted network with unidirectional domains in between, depending on the strength of head-tail attraction (Fig. 3*A*). This effect is irrespective of the length of lamin fibers, *n* (see Supplementary Material Fig. S17-18). While nematic domains of loosely packed chains emerge at *U*_HT_ = 5.0*k*_B_*T*, some bulk aggregation is also observed at *U*_LN_ ≪ *U*_HT_ (i.e. *U*_HT_ = 10.0*k*_B_*T*) (Fig. 3*A*). The intermediate value of lamin-shell attraction strengths tested here, *U*_LN_ = 5.0*k*_B_*T*, is also sufficient to align the chains parallel to one another (Fig. 3*B* and see Supplementary material Fig. S19, S20). However, in these cases, sparse lamin-free regions (see Supplementary material Fig. S20) and bulk accumulation (not shown in Fig. 3) also emerge, particularly if lamin-lamin attraction is stronger than lamin-shell attraction. As the lamin-shell association strength is increased further to *U*_LN_ = 10.0*k*_B_*T*, this trend disappears, and lamin chains cover the entire surface symmetrically, irrespective of the strength of head-tail attraction (Fig. 3*C*).

Overall, our simulations suggest that mutations increasing lamin association with INM can be pivotal for the accumulation of lamin chains near the periphery. Nevertheless, such a pattern can be distorted if the lamin-lamin attraction excessively overcomes lamin-shell attraction. In the next sections, we will discuss how these interactions might govern the effective thickness of the lamin (rod) layer at the periphery and dissociation kinetics of chains from this layer.

### C. Thickness of the lamin layer depends on lamin-lamin association strength and concentration

Previous experimental studies reported thicker nuclear lamina in disease phenotypes due to lamin accumulation towards the periphery [8, 22, 67]. Consistently, our analyses in the previous sections demonstrate that lamin chains can coat the inner surface of the spherical confinement (Fig. 2, 3, and see Supplementary Material Fig. S21, S22). To quantify the thickness of the lamin chain layer in our simulations, we analyze the radial distributions of the lamin chains in the model spherical nucleus. We calculate the positional fluctuations, *dr*, of the center-of-mass-adjusted radial distances for each chain over the simulation time. As the simulation proceeds, if the radial coordinate of a lamin chain fluctuates weakly (i.e., less than the unit size of a bead, *dr* ≤ 0.25) near the shell boundary, the lamin chain is considered immobile and *bound* (Fig. 4*A*). The time-averaged radial position, *R*, of this chain contributes to the thickness of the nuclear lamina via the expression:

**FIG. 4.**
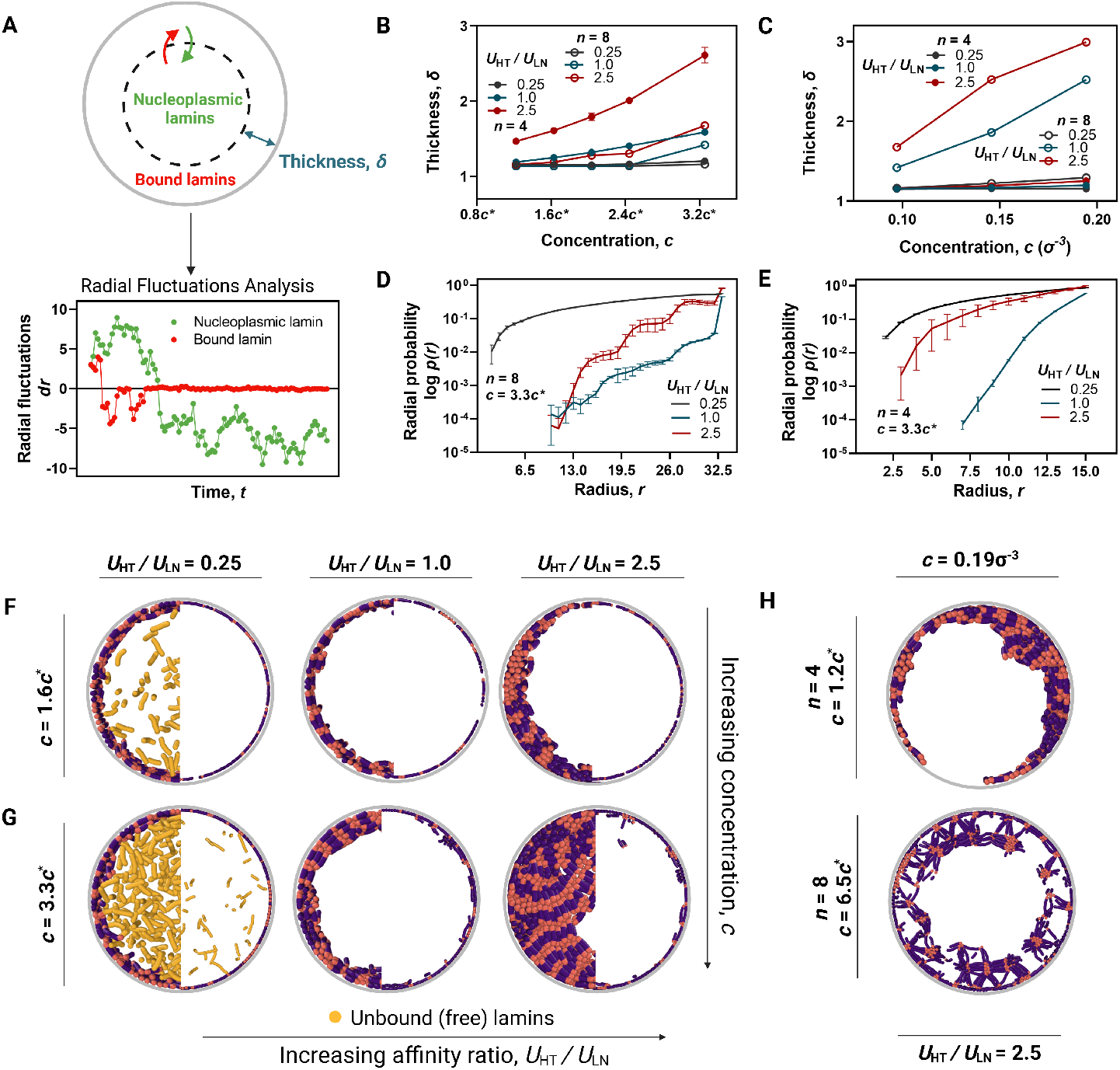
The effect of ratio of affinities, *U*_HT_*/U*_LN_, concentration, *c*, and chain length, *n*, on lamina thickness. (*A*) The methodology used to calculate lamina thickness, *δ*, from simulation trajectories. The schematic shows the fluctuation of radial coordinates for a single *bound* and *unbound* lamin. A lamin is considered *bound* to the shell if its radial coordinate stops fluctuating significantly during a simulation. (*B*) Lamina thickness, *δ*, vs. concentration as a function of *c*\*, *c*, at various affinity ratios and chain lengths. (*C*) Lamina thickness, *δ*, vs. absolute lamin concentration for various *U*_HT_*/U*_LN_. (*D-E*) Radial probability distributions, *p(r)*, as a function of radial distance, *r*. (*F-H*) Representative snapshots of the cross-sections of the nuclei (*F-G*) for various affinity ratios, *U*_HT_*/U*_LN_. (*H*) Representative snapshots for short *n* = 4 (top) and *n* = 8 (bottom) chains.

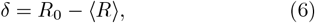

where *R*_0_ is the radius of spherical confinement (Fig. 1*C*), and ⟨*R*⟩ is the average radial position of all lamin chains fulfilling the *bound* condition.

Fig. 4 shows the thickness of the lamin layer, calculated via Eq. 6, as a function of concentration for various ratios of head-tail to lamin-shell affinities, *U*_HT_*/U*_LN_ (see Supplementary Material Fig. S23, S24). In addition to our default chain size (i.e., *n* = 8), calculation results for shorter chains (i.e., *n* = 4) are also shown in Fig. 4. As a general trend, the thickness of the lamina layer increases with increasing *U*_HT_*/U*_LN_, irrespective of chain length (Fig. 4*B, C*).

The thickness of the chain layer at the periphery also increases with increasing concentration (Fig. 4*B, C*). This increase is weak but statistically significant, and consistent with experiments where the peripheral accumulation of mutated lamin A increases with the progression of HGPS (Fig. 4*B*). At the lowest concentration used here (i.e., *c < c*\*), the number of lamin chains in the spherical confinement is only enough to form a single layer with a maximum thickness *δ* ≈ 1.3*σ*, irrespective of the affinity ratio, concentration or chain length *n* (Fig. 4*B, F*). At intermediate (i.e., *c* ≈ *c*\*) to high (i.e., *c >> c*\*) concentrations, the thickness becomes dependent on the ratio of the two competing interactions. For *U*_HT_*/U*_LN_ *<* 1, the thickness has no dependence on concentration (Fig. 4*B, F, G* (first columns)). Notably, at such ratios, there are unbound, freely diffusing chains in bulk (Fig. 4*F, G*). As the ratio *U*_HT_*/U*_LN_ is increased, the effect of concentration becomes more dominant, resulting in the formation of chain multilayers covering the entire surface (Fig. 4*B, G*).

Fig. 4*G* also shows that the thickness increases less significantly for *n* = 8 than *n* = 4 for 0.8*c*\* ≤ *c* ≤ 3.3*c*\* at higher affinity ratios. We hypothesize this could be because the overlap concentration of 8-bead lamins is about 4 times lower than 4-bead lamins, leading to lower absolute concentrations. Consistently, at a higher concentration range (i.e., 6-fold higher than the threshold concentration), 8-bead chains also exhibit thickening at *U*_HT_*/U*_LN_ *>* 1 (Fig. 4*C, H*). Notably, shorter chains accumulate non-uniformly near the periphery, while longer chains form aberrant nodes near the periphery (Fig. 4*H*). To further confirm the partitioning of lamin proteins into *bound* (i.e., those forming the layer) and *unbound* (i.e., free or nucleoplasmic) phases, we also calculate the radial probability distributions, *p*(*r*), by counting the number of lamin chains, *n*(*r*), in a thin slice of the shell’s volume with width Δ*r* = 1.0*σ*:

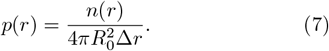

The radial probability distribution for various affinity ratios is consistent with the thickness calculations (Fig. 4*D, E*). At low lamin-lamin interaction strengths with fixed *U*_LN_ = 10.0*k*_B_*T*, specifically *U*_HT_*/U*_LN_ = 0.25, the nuclear lamins are distributed throughout the confinement, co-existing in both the central and peripheral regions of the volume (Fig. 4*D, E*). In contrast, at higher lamin-lamin affinities (i.e., *U*_HT_*/U*_LN_ = 1.0), lamin chains localize at the periphery, leaving the bulk lamin-free (Fig. 4*C, E*). Notably, larger error bars in *U*_HT_*/U*_LN_ *>* 1 case for both chain lengths are consistent with the non-uniform chain distribution and larger thickness of the layer at the periphery (Fig. 4*H*). Collectively, these calculations demonstrate that while weak lamin-shell association can localize some lamin proteins at the periphery, thickening and accumulation at the lamin layer in the model is primarily controlled by head-tail association affinity of the lamin chains, *U*_HT_. This interaction could lead to the segregation of lamins at the nuclear periphery [9] while contributing to the partial thickening of the nuclear lamina in disease.

### D. Extent of lamin chain dissociation depends on head-tail association strength and concentration

One of the hallmarks of HGPS is the suppressed kinetic dissociation of mutant lamins from the nuclear lamina [7]. In accordance with those studies, in some of our simulations, lamin chains unbind from and rebind to the periphery (Fig. 4*F, G* (first column)). In other cases, they are more stably bound to the peripheral aggregates (Fig. 4*H*). To quantify lamin dissociation kinetics thoroughly in our simulations, we calculate the survival fraction of bound lamins as a function of normalized simulation time, *t*, for various concentration and affinity cases (see Methods Section). For simplicity, we choose a fixed lamin-shell attraction strength, *U*_LN_ = 10.0*k*_B_*T*, and vary the head-tail attraction of lamin fibers. However, reducing the strength of lamin-shell attraction from *U*_LN_ = 10.0*k*_B_*T* or reducing chain length (i.e., *n* = 4) does not change the overall lamin dissociation patterns qualitatively as discussed below (Supplementary Material Fig. S25-S29).

Fig. 4*A* shows that, for relatively weaker lamin-lamin affinity (i.e., *U*_HT_*/U*_LN_ = 0.25 with *U*_HT_ = 2.5*k*_B_*T*), as the chain concentration is increased, more free lamins are observed in bulk. Furthermore, increasing the concentrations leads to a more rapid decay in the survival fractions (Fig. 5*A*, right panel): more than half of the bound lamins dissociate within the first half of the simulation time at sufficiently high concentrations (Fig. 5*A*).

**FIG. 5.**
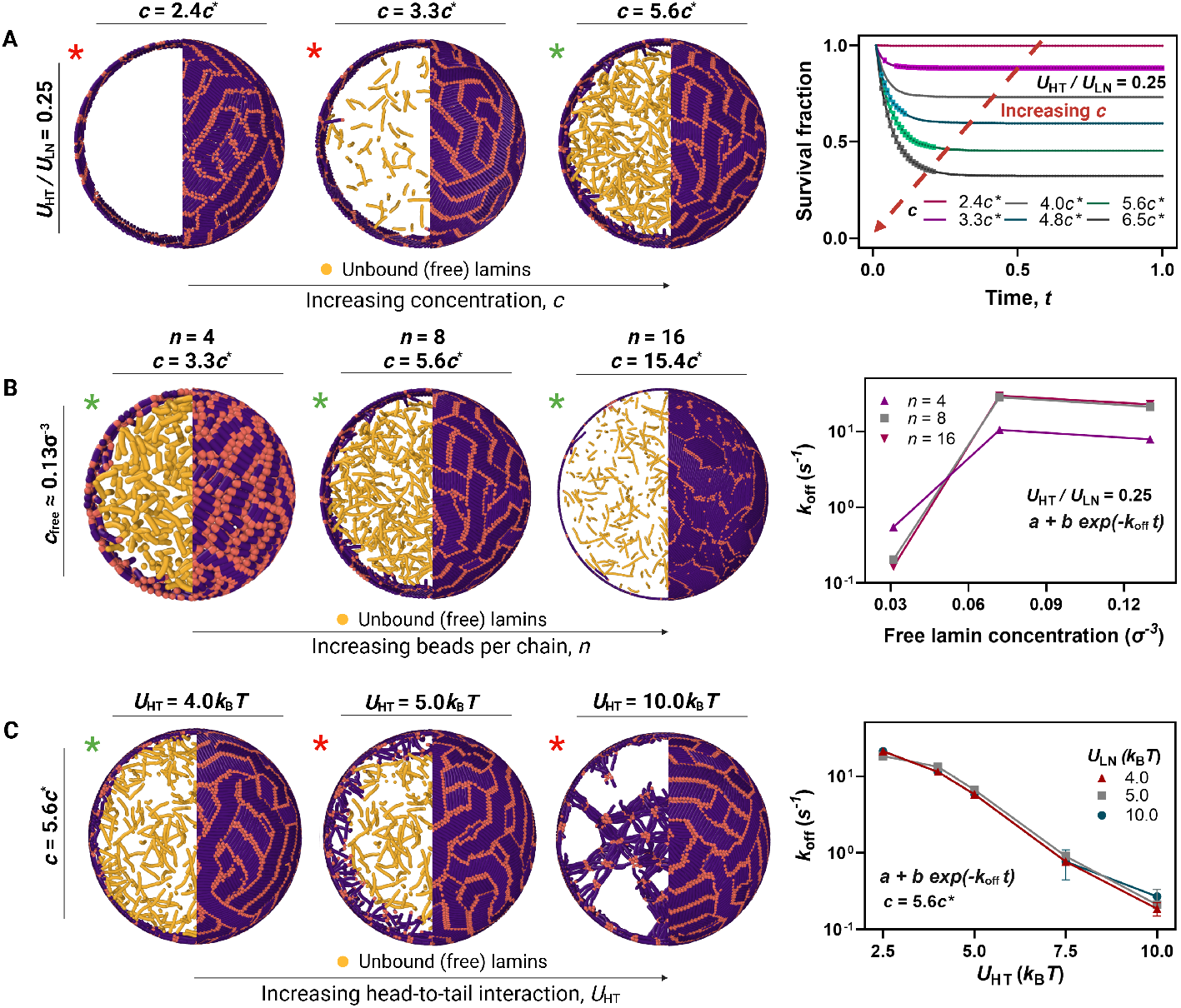
Dependence of lamin dissociation kinetics on concentration, *c*, and head-to-tail association potential to lamin-shell affinity, *U*_HT_. Yellow chains represent *unbound*, free lamin proteins participating in a kinetic exchange. (*A*) Illustrations representing the degree of lamin exchange at three different concentrations with fixed *U*_HT_*/U*_LN_ = 0.25. The panel on the right shows the normalized survival fraction, *n*(*t*)*/n*_0_, of *bound* nuclear lamins over time, *t*. (*B*) Representative snapshots and off-rates as a function of free lamin concentration for *U*_HT_*/U*_LN_ = 0.25 for various lamin chain lengths, *n*. (*C*) Off-rates as a function of head-tail association potential, *U*_HT_, for various *U*_LN_ at a fixed concentration, *c* = 5.6*c*\*.

We next determine the effect of chain length on the kinetic exchange (Fig. 5*B*). Our calculations show that increasing the chain size, *n*, also requires more lamin chains to observe a similar amount of “free” chains in bulk, *c*_free_ (Fig. 5*B*, right panel). Once we extract off rates by fitting survival fraction data to the exponential function in Eq. 5 [68], we observe qualitatively similar patterns in lamin dissociation for comparable *c*_free_ values regardless of chain length. In general, off-rates increase monotonically with *c*_free_, eventually approaching a saturation limit (Fig. 5*B*, right). This dissociation pattern is consistent with the facilitated dissociation (FD) response, where an increase in the concentration of competing species in solution can lead to higher off-rates (i.e., shorter residence times) [60, 65, 68, 69] (see Supplementary Material Fig. S1, S11). Accordingly, if there is an excess concentration of free nucleoplasmic lamins, they could compete with *bound* lamins at the nuclear lamina and speed up their dissociation. Notably, at high concentrations (i.e., *c >> c*\*), the calculated off-rates mostly depend on head-tail association affinity, visually independent of the lamin-shell attraction strength in our simulations (Fig. 5*C*).

Overall, our analyses and calculations imply that increasing lamin head-tail affinity could suppress the dissociation of lamin chains from their aggregates at the periphery, often accompanied by a thicker (nematic) layer of lamin chains at the periphery. This is compatible with experimental observations of suppressed (mutant) lamin dissociation from HGPS lamina [7].

## IV. DISCUSSION

The nuclear lamina can adapt to changing physiochemical and mechanical conditions by altering the concentration and solvation properties of its main constituents, lamin proteins. In this study, we model supramolecular structures of lamin proteins (i.e., fibers) as stiff, short polymer chains. This over-simplification allows us to explore the physical principles and relative interactions that can lead to experimentally observed morphological and kinetic changes in lamin’s (particularly A-type lamins) phase behavior associated with laminopathies such as accelerated aging (i.e., Progeria). In our simulations, the spherical, rigid confinement represents the nuclear void bounded by the inner nuclear membrane (INM), which can interact with lamin proteins. The lamin model attempts to mimic the complex head-tail association of lamin fibers by incorporating asymmetric interactions between chain ends. This simple model reproduces the nematic phase formation, nuclear lamina thickening, and suppressed dissociation kinetics of mutant lamin proteins observed in diseased cell nuclei [7, 8, 39, 43]. It also emulates certain properties of the nuclear lamina in healthy cells, all by exploiting lamin-INM interactions, lamin-lamin association affinities, and lamin concentration. Below, we will further discuss the potential implications of our calculations in context of the structural biology of cell nuclei in health and disease.

### A. Alterations in lamin localization to the nuclear periphery can lead to structural anomalies in nuclear lamina

Once cell division initiates, the nuclear lamina disassembles via a hyperphosphorylation mechanism and later re-assembles around the replicated chromosomes of the sister cells [11, 12]. A prerequisite for this re-assembly is the localization of some, if not all, lamin proteins (mostly lamin A/C since lamin B is often attached to INM) or their supramolecular structures to the nuclear periphery. This way, lamins can associate with one another and/or with the INM to form a two-dimensional (2D) meshwork underneath the INM. Importantly, this self-assembled and dynamic meshwork must provide mechanical support to the nucleus while ensuring nuclear shape stability in a healthy cell.

In our simulations, the localization of rod-like lamin chains results from the competition between the lamin self-affinity and their tendency to bind to the periphery of the spherical shell (i.e., the INM) (Fig. 4). Lamin-shell affinity comparable to the lamin-lamin (head-to-tail) association strength ensures the formation of a uniformly distributed lamin layer at the periphery (Fig. 4*G*), where lamin chains organize more an isotopic, network-like fashion, consistent with the lamin organization in healthy cells (Fig. 4*A, B*).

The nuclear lamina becomes thicker and mechanically more rigid in HGPS due to the excess aggregation of mutant lamin A, progerin, at the nuclear lamina [7, 41, 42], causing decreasing morphological adaptability in the disease. From a molecular perspective, progerin remains farnesylated after post-translational modifications. Farnesylation promotes progerin’s membrane association and enhances its thermodynamic stability compared to *wt* lamin A [70, 71]. Notably, increased hydrophobic interactions have also been reported between the tail domain of progerin and INM proteins, such as SUN1 and Emerin [28]. These experiments collectively suggest that truncated lamin A might have a stronger tendency to associate with the INM compared to *wt* - lamin A. Our simulations support this, showing that strong lamin-shell attraction (i.e., strong binding to the INM) can lead to strong aggregation of chains towards the periphery (Fig. 2, 3). Further, under a finite lamin-lamin association strength, the strong peripheral localization gives rise to nematic-phase domains in calculations, which were realized experimentally both *in vitro* [36, 72] and *in vivo* [7]. Noteworthy, long semi-flexible chains were shown to form global nematic phases (see Supplementary Material Fig. S2, S6, S7) [48–51], unlike the finite-size domains observed in our study, further supporting the notion that in Progeria, lamin fibers might be shorter than their healthy counterparts.

Moreover, the formation of nematic domains may require stronger hydrophobic interactions between lamins at the nuclear surface, which is often the case in liquid crystals [47, 73–76]. Studies on the crystal structure of progeria mutant *S*143*F* protein suggested the formation of X-shaped disulfide bonds in the central rod domains of progerin [18]. This implies that hydrophobic interactions can play a role in altering the morphology of the lamina meshwork [18, 71], possibly bringing cysteine residues of progerin molecules close to one another. In light of our results, which treat the lamin proteins as hydrophobic (except their head and tail domains), the enhanced hydrophobic interactions in HGPS could also contribute to forming nematic phase domains at the nuclear lamina.

### B. Inter-lamin affinity can dictate nuclear lamina thickening

In addition to the formation of nematic domains, nuclear lamina thickening also serves as a key disease hallmark along with abnormal nuclear shape [6, 8, 24–29, 67]. Accordingly, as progerin concentration increases parallel to the disease progression, the nuclear lamina thickens as toxic mutant proteins accumulate at the periphery [8, 43]. Such aggregation can be promoted by an increase in the effective affinity of mutant lamins towards each other or the INM. This view is also consistent with the observed decrease in exchange rates of progerin [7], indicating an increase in the interaction energies between mutant proteins and their molecular environment. In line with this, our simulations suggest that in HGPS, the thickening of the nuclear lamina near the periphery can be the result of an interplay between lamin localization and increasing lamin-lamin association (e.g., head-tail) as the nuclear concentration of mutant lamins increases (Fig. 4). In fact, relatively higher head-tail association strength readily results in thicker chain layer and chain aggregation in simulations, suggesting a dominant effect for inter-lamin interactions.

Consistent with this view, permanent farnesylation of progerin enhances its binding properties to the INM compared to *wt* - lamin A [71], as discussed earlier. However, the farnesyl group remains sequestered within a hydrophobic region in the tail domain unless physiological concentrations of divalent ions, such as *Ca*^2+^, are present at the nuclear periphery [67]. Without calcium ions, the farnesyl group only acts as a reversible crosslink, temporarily associating with membrane proteins, but can still dissociate into the nucleoplasm, a characteristic of healthy nuclei [19, 71]. Calcium can induce a conformational change in progerin, opening up the tail domain’s Ig-like fold and exposing the farnesyl group for molecular interactions. This exposure can facilitate stronger inter-molecular interactions with other lamins (e.g., progerin and other unaltered lamins) and the INM [26, 67]. These experimental observations are consistent with our MD simulations, where nuclear lamins become constrained to the INM when both head-tail association and lamin-shell association strengths are high (i.e., *U*_HT_ ≥ *U*_LN_ *>* 1*k*_*B*_*T*) (Fig. 4), suggesting an intricate interplay of intermolecular interactions promoting lamina thickening at the nuclear periphery.

### C. Dissociation kinetics of lamins from nuclear lamina can have a concentration dependence

Photobleaching experiments suggest that peripherally bound lamins could exchange with their nucleoplasmic counterparts in healthy cells [7]. One possible mechanism by which free lamins can influence the kinetics of lamin exchange between those in the nuclear lamina and nucleoplasm (free) in health could be facilitated dissociation (FD). FD is a concentration-dependent dissociation mechanism, originally discovered in protein-DNA interactions [60, 68, 69, 77]. FD causes an increased exchange (dissociation) rate for a ligand attached to its binding site when free ligands are present in the solution. The mechanism is based on the molecular-scale competition between the two ligands at the binding site (such as a third lamin) (see Supplementary Material Fig. S1, S11, S25-S29). Nucleoplasmic lamin proteins can likely compete with their bound counterparts in the nuclear lamina, leading to rapidly decaying signals observed in the experiments. Our simulations demonstrate such a concentration-dependent mechanism for lamin chains of varying sizes (Fig. 5). In contrast, the exchange rate drops in HGPS [7], possibly due to increasing inter-molecular interactions between the mutant protein and the nuclear lamina, and depletion of lamin from nucleoplasm. Consistently, our calculations show that increased lamin-lamin affinity reduces dissociation rates while promoting further lamin accommodation near the periphery and eventually affecting the thickness of the chain layer (Fig. 4, 5). Furthermore, unlike HGPS cells, the accumulation of lamin proteins (specifically *wt* - lamin A) increases nuclear stability in some healthy cell lines [41, 42], and even correlates with decreased DNA double-strand breaks [42]. Thus, based on the available literature and our simulations, we speculate that lamin unbinding kinetics from the nuclear lamina could also have structural consequences, leading to either a highly adaptive, mechanically soft lamina or a highly stable but mechanically brittle one.

In summary, we employed a coarse-grained polymer model of lamin supramolecular structures to investigate the key physical interactions that can drive the abnormal self-assembly of lamin proteins in the formation of the nuclear lamina, with a particular focus on HGPS (Progeria syndrome). Our findings suggest that enhanced inter-molecular interactions among mutant lamins can lead to specific disease hallmarks in the nuclear lamina structure. The simulations reveal an interplay between lamin-lamin and lamin-INM interactions in a concentration-dependent manner, demonstrating how this interplay can reduce lamin exchange rates, cause lamina thickening, and promote the formation of nematic microdomains, as observed in HGPS.

### D. Limitations of the lamin model

Although our coarse-grained model can recapitulate several experimental phenotypes observed in Progeria, namely: i) phase transition of the lamina from an isotropic to nematic organization; ii) lamina thickening; and iii) lamin dissociation kinetics between the lamina and interior, a conclusive understanding of lamina assembly and structure in disease is far from complete. We assume a uniformly attractive spherical confinement to model the INM. In reality, the INM exhibits spatially heterogeneous mechanical and biochemical properties [17, 22, 44, 53]. This spherical confinement is also rigid, unlike the eukaryotic nucleus, which undergoes significant shape distortions in diseases like Progeria. Furthermore, given that the disease is caused by a *de novo* point mutation in the *LMNA* gene that encodes for human lamin A/C [20], our model focuses on a single lamin type only. On the contrary, the lamina meshwork in healthy cells is composed of heterogeneous, network-like arrangements of both A- and B-type lamins [23]. Our model also oversimplifies the assembly of lamin supramolecular structures such as protofilaments and other higher-order structures by modeling lamin chains as a monodisperse solution of semi-flexible rods, ignoring dimers or monomeric lamins. The model also does not include chromatin, which can affect lamin organization or *vice versa*. It simplifies overall parameters that might affect the nuclear lamina organization into concentration, inter-lamin, and lamin-shell interactions and ignores any specific interactions that can play key roles in disease.

Future studies could benefit from more sophisticated models to better understand these experimental features such distorted nuclear morphology and mechanics in HGPS [8, 29, 38–40]. For instance, replacing spherical confinement with elastic shell models [78, 79] filled with chromatin [54] could provide deeper insights into nuclear shape alterations associated with the disease.

## Supporting information

Supplementary Data for Phase Behavior and Dissociation Kinetics of Lamins in a Polymer Model of Progeria

## V. ACKNOWLEDGMENTS

AE acknowledges E.J. Banigan for his suggestions on disseminating this work. This research was supported by TUBITAK, The Scientific and Technological Research Council of Turkey [1001 Grant No. 122F309] and The National Science Center, EU’s H2020 Programme, and MSCA Grant Agreement No. 945339 Poland [Grant Polonez Bis No. 2021/43/P/ST3/01833].

