## Supplementary Data for Phase Behavior and Dissociation Kinetics of Lamins in a Polymer Model of Progeria for "Phase Behavior and Dissociation Kinetics of Lamins in a Polymer Model of Progeria"

### Supplementary Data and Figures for Phase Behavior and Dissociation of Lamins in a Polymer Model of Progeria

#### A. Molecular Dynamics (MD) Simulation Parameters

In our MD simulations, each lamin fiber is modeled using coarse-grained "Kremer-Grest (KG)" bead-spring chains in implicit solvent. Lamin fibers were confined within a spherical confinement of radius, $R_0$. $R_0$ is fixed to $34\sigma$ for 8-bead coarse-grained lamin fibers. All lamin proteins are made of identical beads of size $b = 1\sigma$, where $\sigma$ represents the unit length in the MD simulations. The monomeric LJ mass for all beads is set as $m = 1$.

All non-bonded pairwise interactions were defined with Lennard-Jones (LJ) potential. The LJ potential was shifted and truncated as shown below:

$$V_{\mathrm{LJ}}(r) = \begin{cases} 4u\left[\left(\frac{\sigma}{r}\right)^{12} - \left(\frac{\sigma}{r}\right)^{6} + v_s\right] & r \le r_c \\ 0 & r > r_c \end{cases} \tag{1}$$

The default cut-off distance is $r_c = 2^{\frac{1}{6}}\sigma$, and the shift factor as $v_s = \frac{1}{4}$. The strength of the repulsive interaction is $u = k_{\mathrm{B}}T$, unless otherwise stated. $k_{\mathrm{B}}$ is the Boltzmann constant, and $T$ is the absolute temperature. For all attractive potentials defined in the simulation i.e. spherical confinement, and between lamin fibers, the cut-off distance is set as $r_c = 2.5\sigma$ with a shift factor of $v_s = 0$.

Two main types of intermolecular interactions were investigated: lamin-shell interaction, $U_{\mathrm{LN}}$, and lamin-lamin association, $U_{\mathrm{HT}}$. The attraction strength combinations of these two investigated in our simulations and discussed in the main text are defined in Table I. $U_{\mathrm{LN}}$ $= 10.0k_{\mathrm{B}}T$ cases are discussed in most detail in the main text (Fig. 2, 5), while the others are shown below.

| $U_{\mathrm{LN}}$ ($k_{\mathrm{B}}T$) | $U_{\mathrm{HT}}$ ($k_{\mathrm{B}}T$) | | | | |
|---|---|---|---|---|---|
| 2.5 | 2.5 | 4.0 | 5.0 | 7.5 | 10.0 |
| 4.0 | 2.5 | 4.0 | 5.0 | 7.5 | 10.0 |
| 5.0 | 2.5 | 4.0 | 5.0 | 7.5 | 10.0 |
| 10.0 | 2.5 | 4.0 | 5.0 | 7.5 | 10.0 |

TABLE I: The interaction energy combinations of lamin-nucleus interaction, $U_{\mathrm{LN}}$, and lamin-lamin attraction, $U_{\mathrm{HT}}$, that are investigated in our simulations.

All bonded interactions between adjacent beads in each lamin fiber are assigned with a non-extensible FENE potential.

$$V_{\mathrm{Bond}}(r) = -0.5kr^2 \ln\left[1 - \left(\frac{r}{r_0}\right)^2\right] \tag{2}$$

where the bond energy was set to the default value, i.e. $k = 30.0k_{\mathrm{B}}T/\sigma^2$. The maximum bond distance was set to $r_0 = 1.5\sigma$. Each lamin fiber is made semi-flexible in nature via a harmonic angle potential:

$$V_{\mathrm{Bend}}(\theta) = k_\theta(\theta - \theta_0) \tag{3}$$

where the energy is set to $k_\Theta = 10.0k_{\mathrm{B}}T/rad^2$, and the reference angle as $\theta_0 = \pi$.

Unless stated otherwise, the time step in the simulation was $\Delta t = 0.005\tau$, where $\tau$ has units of time. To promote Brownian dynamics in our simulations, we used the Langevin thermostat keeping the temperature constant. Since we use the second half of the simulations for data analyses, our simulations remain unaffected by the initial configuration. During the simulation, all of the lamins remain within the spherical confinement. Each simulation was run for $10^6$ MD time steps, and the dumping coefficient was set to $5000\tau$ unless mentioned otherwise. Periodic boundary conditions were used in all directions, and our simulation box was $50 \times 50 \times 50\sigma^3$. All simulations were carried out in the LAMMPS MD simulations package.

##### B. Data Analysis Details

*Lamin concentration, c*

To characterize the concentration regime of the lamins, we convert the number of lamin rods, $N$, to concentration, $c$ (see main text Table I.):

$$c = \frac{Nn}{V}. \tag{4}$$

$$\text{where} \quad n = 8, \quad V = \frac{4}{3}\pi R_0^3, \quad R_0 = 34\sigma.$$

We also define the overlap parameter, $P$ (i.e. the ratio of the concentration to overlap concentration).

$$P = \frac{c}{c^*} = N\frac{v}{V}. \tag{5}$$

We normalize all concentrations in terms of the overlap concentration, $c^*$, and their ratio is known as the overlap parameter, $P$. Lamin concentration, $c$, is discussed in terms of $c^*$, and as absolute concentration in $\sigma^{-3}$. For the 8-bead lamin rods, the overlap concentration, $c^*$ is as calculated follows:

$$\begin{aligned} c^* = \frac{n}{v} &= \frac{8}{\frac{4}{3}\pi(4\sigma)^3} \\ &= 0.02984155183 \approx 0.03\sigma^{-3} \end{aligned} \tag{6}$$

$$\text{where} \quad n = 8, \qquad v = \frac{4}{3}\pi r^3, \quad r = \frac{8\sigma}{2} = 4\sigma$$

Since overlap concentration treats each lamin chain as a single bead with an excluded volume, 4-bead lamin rods have a larger overlap concentration such that:

$$\begin{aligned} c^* = \frac{n}{v} &= \frac{4}{\frac{4}{3}\pi(2\sigma)^3} \\ &= 0.1193662073 \approx 0.1\sigma^{-3} \end{aligned} \tag{7}$$

$$\text{where} \quad n = 4, \qquad v = \frac{4}{3}\pi r^3, \quad r = \frac{4\sigma}{2} = 2\sigma$$

Thus, the overlap concentration, $c^*$, decreases for longer lamin fibers, given that they have a higher $v$.

*Radial Fluctuation Analysis*

To calculate the thickness of the nuclear lamina (NL), we identify lamin proteins that associate with the nuclear periphery irreversibly. In other words, lamin fibers whose radial coordinate stops fluctuating significantly over time are associated irreversibly with the nuclear periphery and are considered *bound*. To find such lamins, we take the derivative of the radial coordinate over time, *dr*. Lamin fibers that satisfy the following condition are considered bound to be part of the lamina thickness:

$$dr \leq 0.25 \tag{8}$$

The time average of the radial coordinates of these bound lamins, $\langle R \rangle$, is used to determine thickness, $\delta$.

$$\delta = R_0 - \langle R \rangle \tag{9}$$

*Radius of Gyration*

To further support the trends we observe for radial probability and thickness, we also calculate an effective thickness, $\delta'$ from average radius of gyration, $R_g$ over all lamin fibers and time steps and define $\delta'$ as:

$$\delta' = R_0 - \langle R_g \rangle \tag{10}$$

*Quantifiying lamin exchange kinetics: free lamin concentration,* $c_{\text{free}}$

As described in the main text, we quantify lamin exchange kinetics in our simulations by counting the number of *bound*, surviving lamins, $n(t)$ at each time step, for the latter half of the simulations. We also calculate time-averaged concentration of *unbound*, free lamins, $c_{\text{free}}$ as follows:

$$c_{\text{free}} = \frac{n_0 - \langle n(t) \rangle}{V} \tag{11}$$

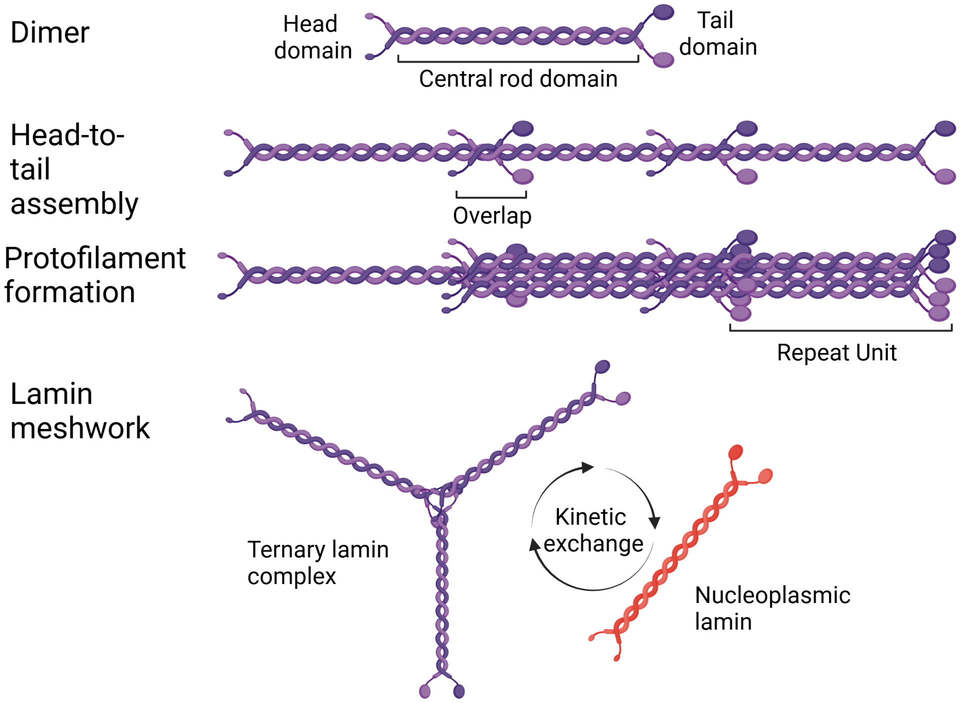


FIG. S1: Potential stages of lamin organization leading to the formation of the nuclear lamina meshwork that lines the inner nuclear membrane in eukaryotic nuclei.

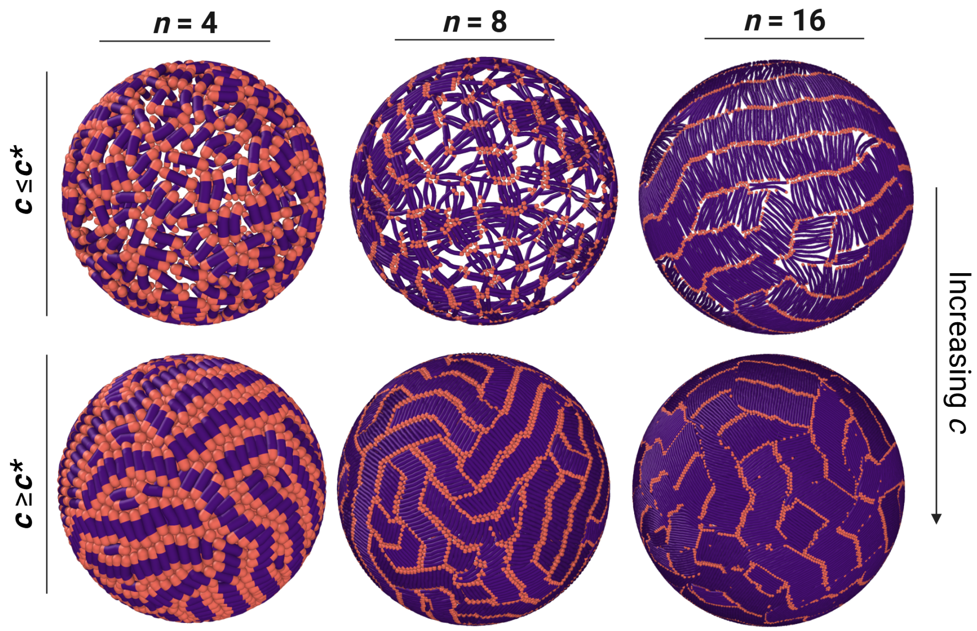


FIG. S2: Isotropic to nematic phase transition of lamin proteins can be observed irrespective of the number of beads constituting each lamin fiber, $n$, provided that the lamin fibers are semi-flexible. Representative snapshots of the spherical confinement when $U_{\mathrm{HT}} = U_{\mathrm{LN}} = 10.0 k_{\mathrm{B}}T$. Increasing concentration, i.e. when $c \geq c^{*}$, results in a phase transition when all interaction strengths are high.

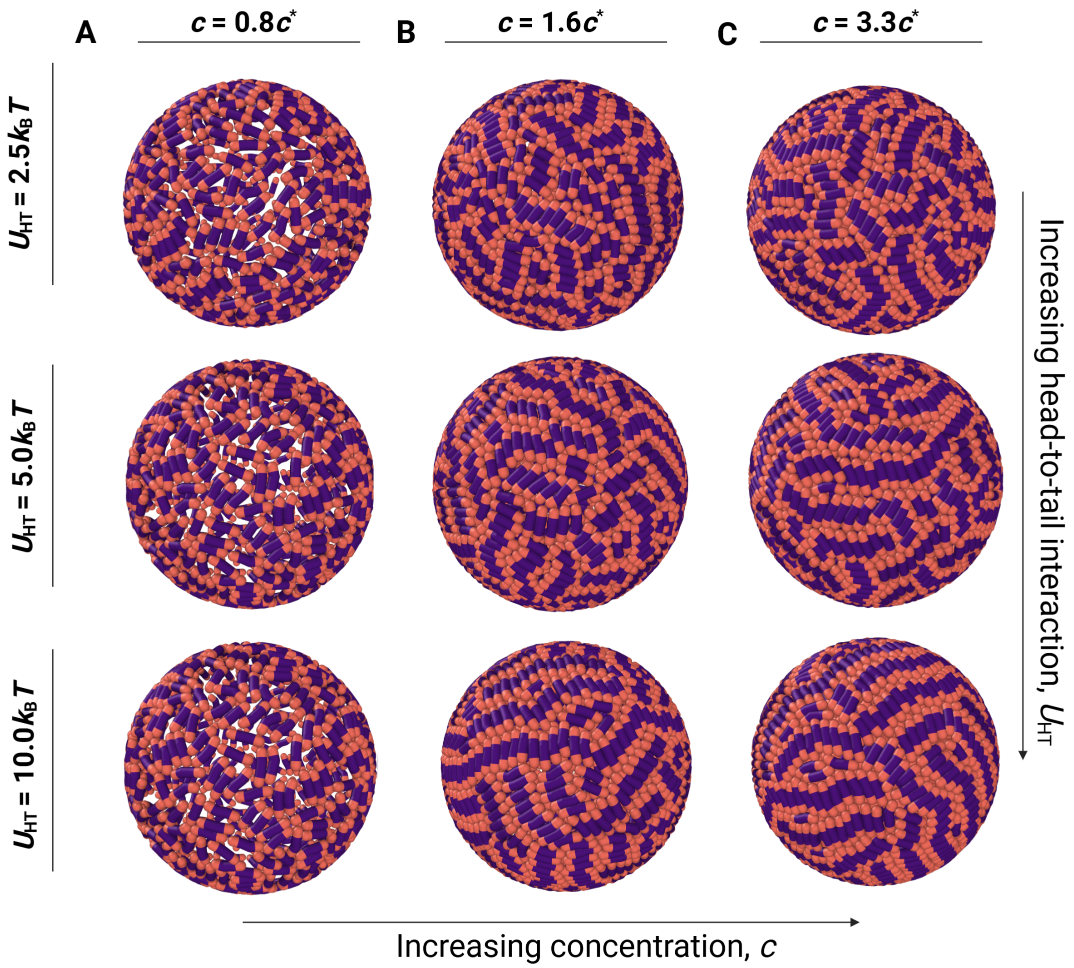


FIG. S3: Representative snapshots for 4-bead lamin fibers at fixed $U_{\mathrm{LN}} = 10.0 k_{\mathrm{B}}T$ when $U_{\mathrm{HT}} = 2.5$, 5.0, and $10.0 k_{\mathrm{B}}T$ for 3 concentrations, **A)** $0.8c^*$, **B)** $1.6c^*$, **C)** $3.3c^*$. The radius of the spherical confinement is halved to $R_0 = 17\sigma$ to conserve concentrations, $c$, used in 8-bead system. Organized microdomains are observed at high concentration irrespective of head-to-tail association potential, $U_{\mathrm{HT}}$.

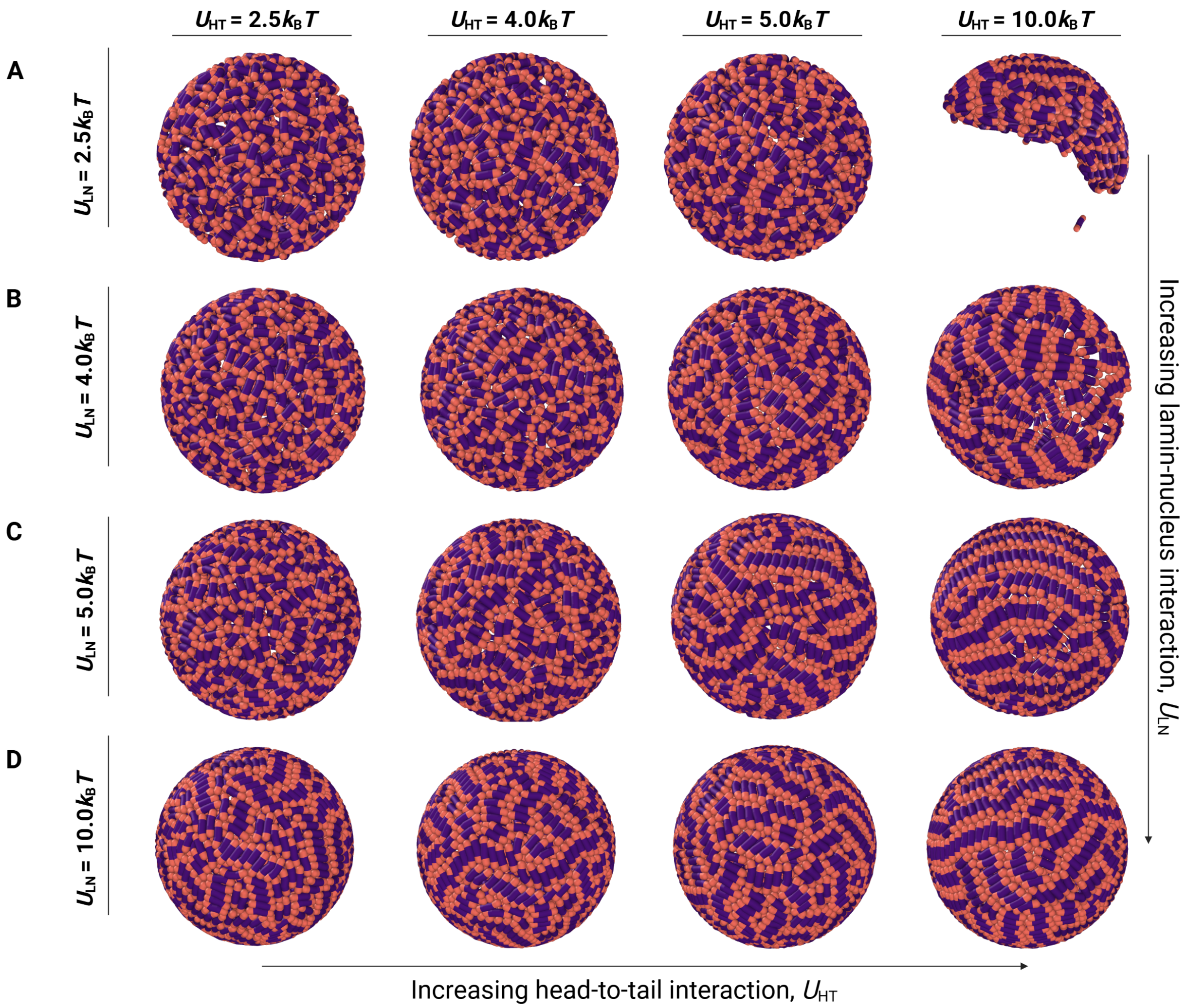


FIG. S4: Isotropic, network-like arrangement of lamin proteins occurs when $c < 2c^*$ for the 4-bead lamin fiber system. Snapshots for $c = 1.6c^*$ case showing the interplay of $U_{\mathrm{HT}}$ and $U_{\mathrm{LN}}$, where $U_{\mathrm{LN}} =$ **A)** 2.5, **B)** 4.0, **C)** 5.0, **D)** $10.0k_{\mathrm{B}}T$.

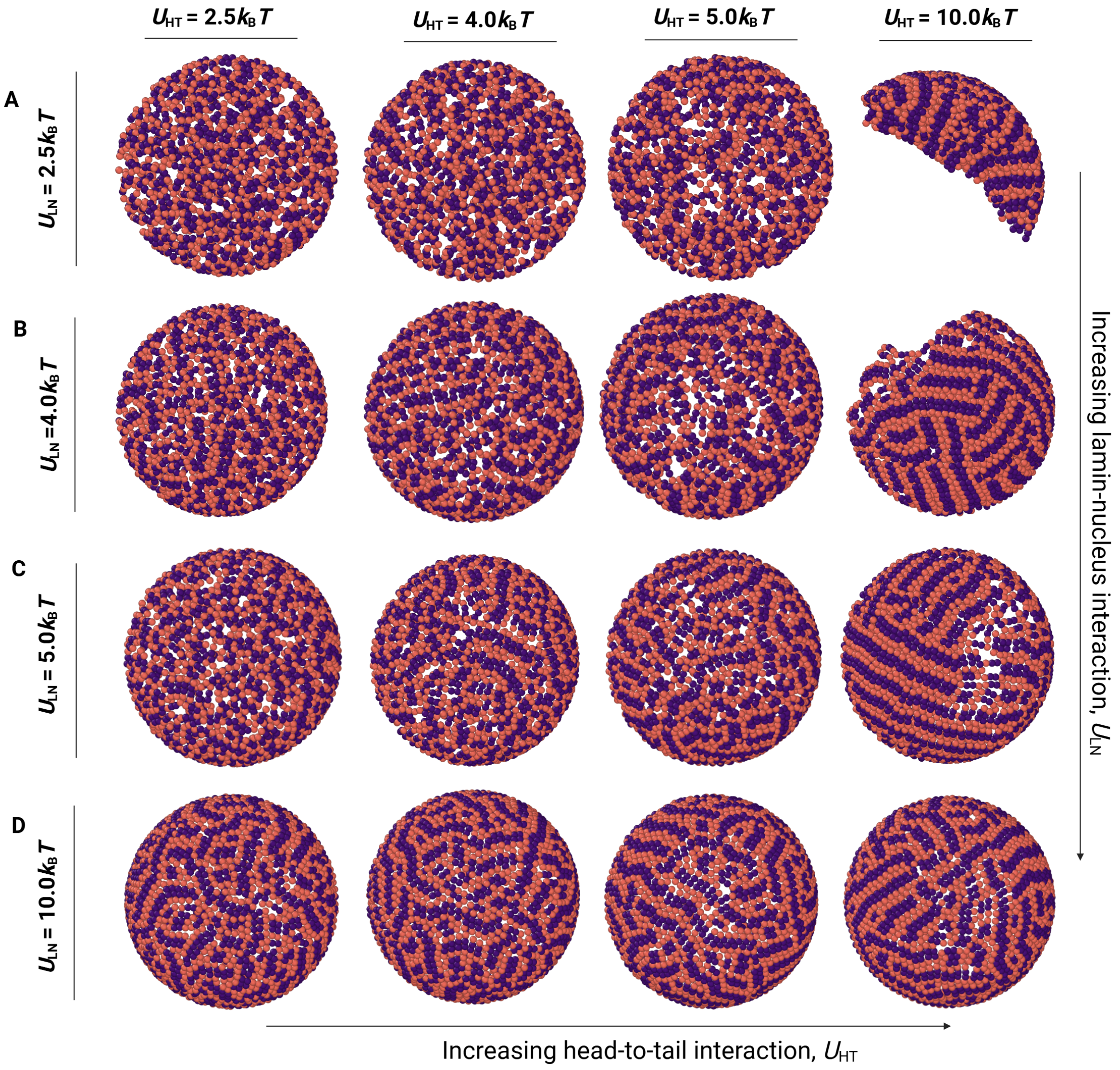


FIG. S5: Organized domains are observed largely irrespective of the head-to-tail association potential of lamin proteins at higher concentrations. Snapshots for $c \geq 2c^*$ case i.e. $c = 3.3c^*$ showing the effect of $U_{\rm HT}$ for $U_{\rm LN} =$ **A)** 2.5, **B)** 4.0, **C)** 5.0, and **D)** 10.0$k_{\rm B}T$.

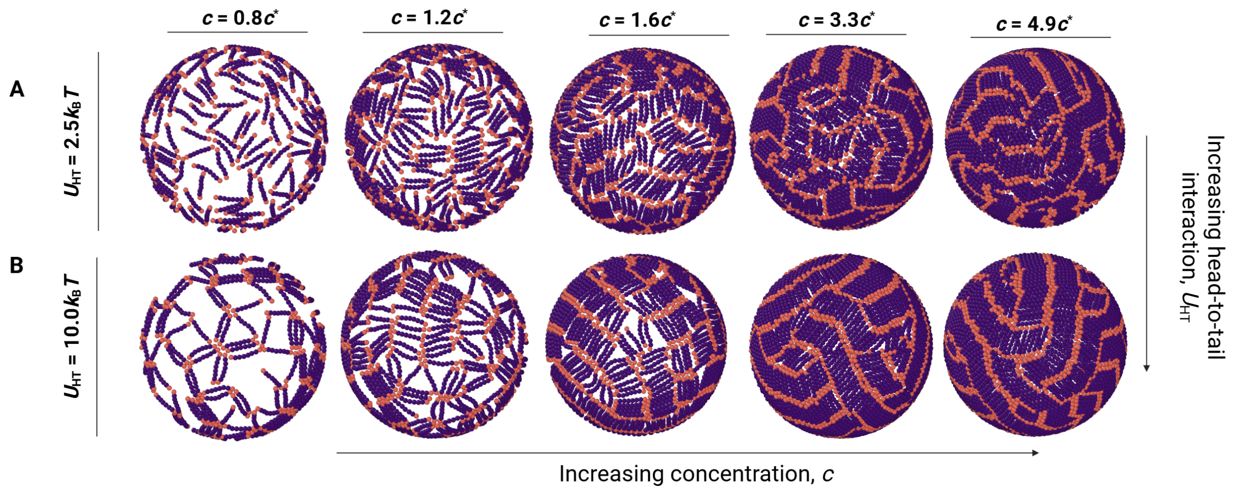


FIG. S6: Microdomain organization of lamin proteins in our system is independent of the radius of the spherical confinement, $R_0$. Representative snapshots of the exterior cross-sections for 8-bead lamin fibers at fixed $U_{\mathrm{LN}} = 10.0k_{\mathrm{B}}T$ when $U_{\mathrm{HT}} =$ **A)** $2.5k_{\mathrm{B}}T$, and **B)** $10.0k_{\mathrm{B}}T$. The radius of the spherical confinement is set to $R_0 = 21.4\sigma$.

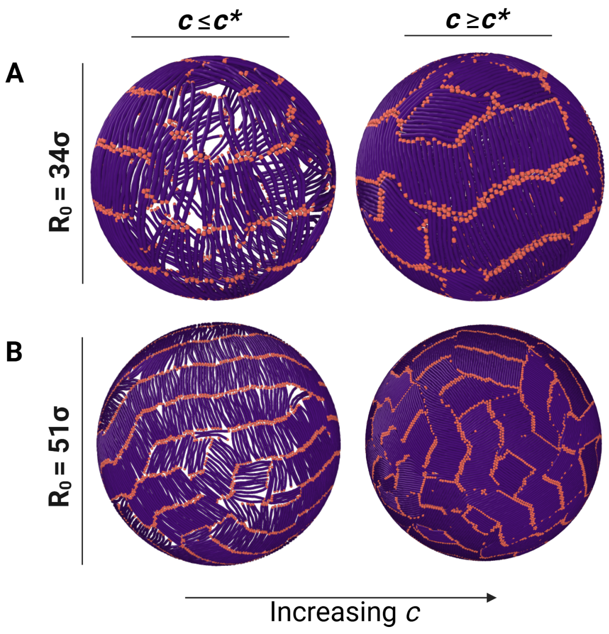


FIG. S7: Increasing concentration results in unidirectional organization of the nematic microdomains for longer lamin proteins when the radius of the spherical confinement, $R_0$, is reduced. Representative snapshots of the spherical confinement when $U_{\mathrm{HT}} = U_{\mathrm{LN}} = 10.0k_{\mathrm{B}}T$ below and above the overlap concentration, $c^*$ for **A)** $R_0 = 34\sigma$ (same as main text), and **B)** $R_0 = 51\sigma$.

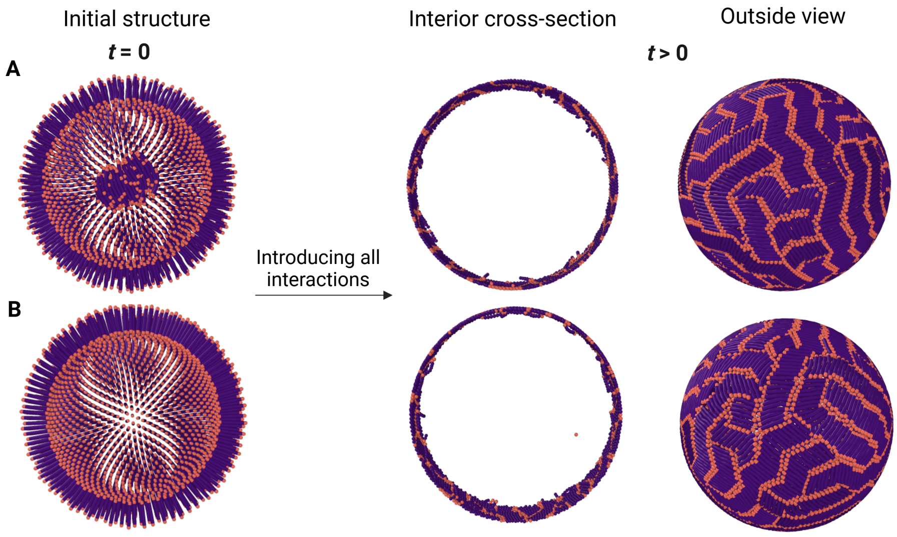


FIG. S8: Our initial configuration does not affect simulation results. Concentration is fixed to $c = 3.3c^*$ for $n = 8$ bead lamins. Both head-tail association potential and lamin shell attraction are also fixed; $U_{\rm HT} = U_{\rm LN} = 10.0 k_{\rm B}T$. **A)** 25% of the lamin fibers are placed quasi-randomly close to the center of the spherical confinement. **B)** All lamins are placed in a spherical manner close to the shell. Similar nuclear lamina thickening and nematic phase formation is observed in both cases irrespective of their initial arrangement.

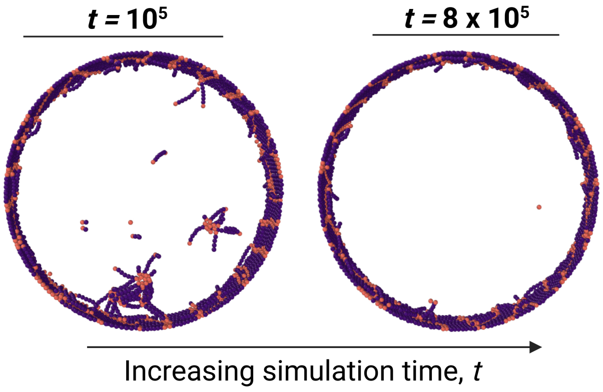


FIG. S9: $10\sigma$ wide slices representing formation and disappearance of aggregates of 8-bead lamin chains in the bulk. **A)** Aggregates form at the center for 8-bead lamin chains, $n = 8$, during the aggregation of lamin chains near the spherical boundary at $t > 0$. **B)** Small aggregates disappear and lamin chains form a uniform lamina at the periphery for a longer simulation time i.e. when the system reaches equillibrium

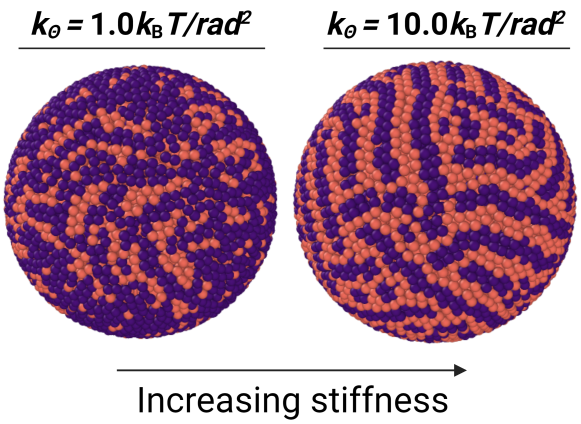


FIG. S10: The effect of stiffness on the microdomain organization of the nuclear lamina for $n = 4$ bead lamins. The concentration is set to $c = 3.3c^*$, and interaction strengths to $U_{\rm HT}$ $= U_{\rm LN} = 10.0 k_{\rm B} T$. The angle potential, Eqn. (3), is increased from fully flexible $(1.0 k_{\rm B} T/rad^2)$ to semi-flexible rod-like $(10.0 k_{\rm B} T/rad^2)$. Lamins need to be semi-flexible to form nematic phases at higher concentrations.

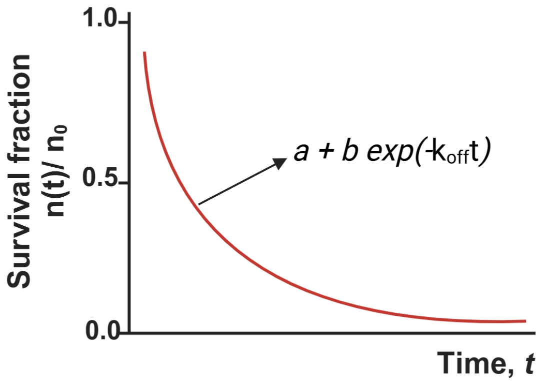


FIG. S11: Sample lamin dissociation curve of normalized survival fraction of lamin proteins, $n(t)/n_0$, vs. normalized simulation time, $t$. Dissociation curves are used to extract off-rates, $k_{\rm off}$.

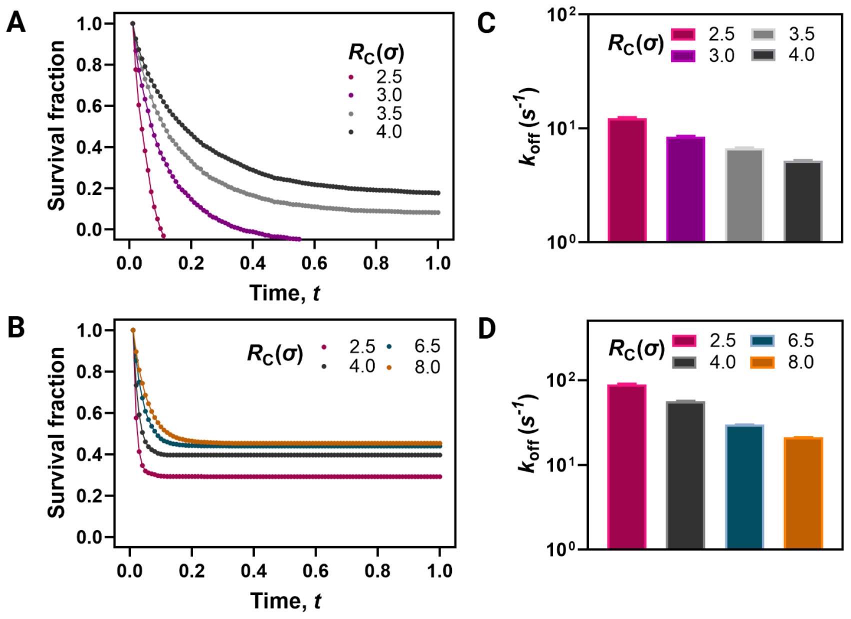


FIG. S12: Benchmarking of the survival fraction and dissociation rates at various threshold distances, $R_{\mathrm{C}}$. Survival fraction at various threshold distances when $U_{\mathrm{HT}} = 2.5k_{\mathrm{B}}T$, and $U_{\mathrm{LN}} = 10.0k_{\mathrm{B}}T$ for **A)** $n = 4$ and **B)** $n = 8$ bead lamins. Dissociation rates of lamin unbinding at different $R_{\mathrm{C}}$ for **C** $n = 4$ and **D)** $n = 8$ bead lamins. $R_{\mathrm{C}} = 4.0\sigma$ for $n = 4$, and $R_{\mathrm{C}} = 8.0\sigma$ for $n = 8$ i.e. respective lengths lamin fibers, were chosen as the upper bound after several trials.

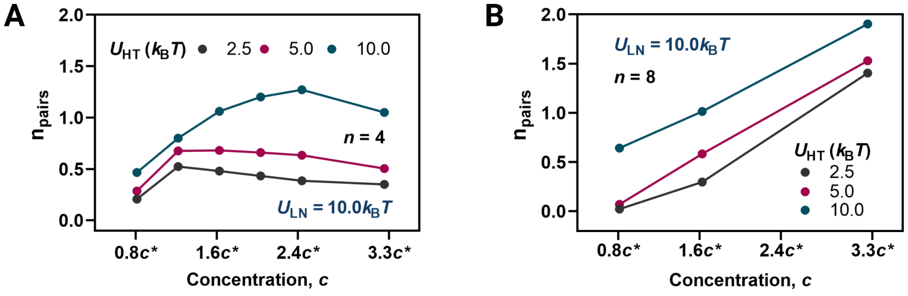


FIG. S13: Normalized number of aligned pairs, $n_{\mathrm{pairs}}$, as a function of concentration, $c$ at high lamin-shell interaction strength, $U_{\mathrm{LN}} = 10.0\ k_{\mathrm{B}}T$ for **A)** $n = 4$ and **B)** $n = 8$-bead lamin fibers. $0 \leq n_{\mathrm{pairs}} \leq 2$ given that the total number of fiber pairs $= 2 \cdot$ total number of lamin fibers.

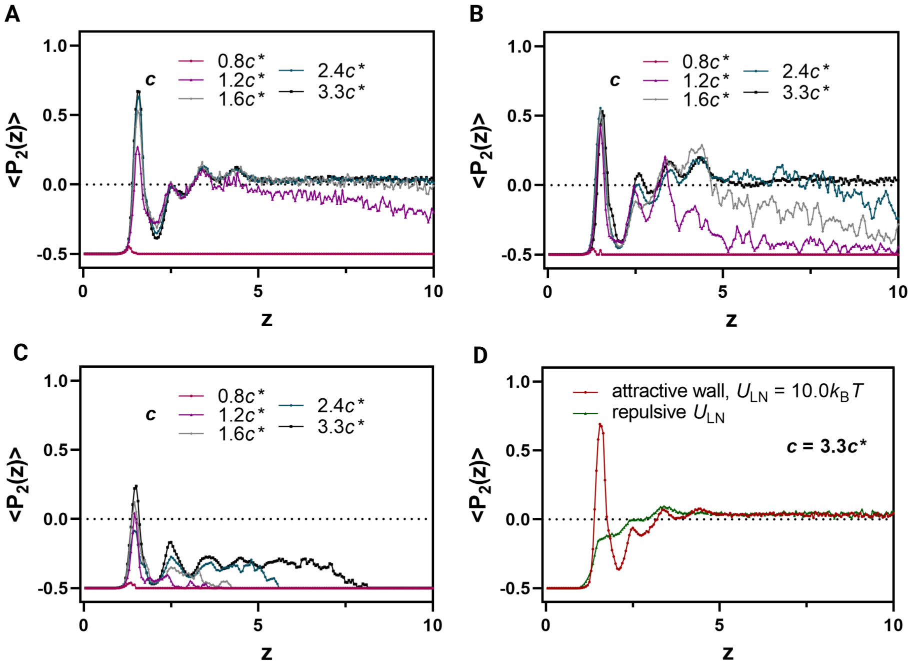


FIG. S14: Second Legendre polynomial, $P_2(z)$, for different values of $U_{\mathrm{HT}}$: **(A)** 2.5, **(B)** 5.0, and **(C)** 10.0 $k_\mathrm{B}T$, with a fixed $U_{\mathrm{LN}} = 10.0\ k_\mathrm{B}T$. **(D)** Control cases showing the effect of $U_{\mathrm{LN}}$ alone on $P_2(z)$. $P_2(z)$ quantifies the orientation of bond vectors $b_i$ at a distance $z$ from the spherical surface, forming an angle $\theta(z)$ with the surface normal vector $n_i$:

$$P_2(z) = \frac{3\cos^2\theta(z) - 1}{2}. \tag{13}$$

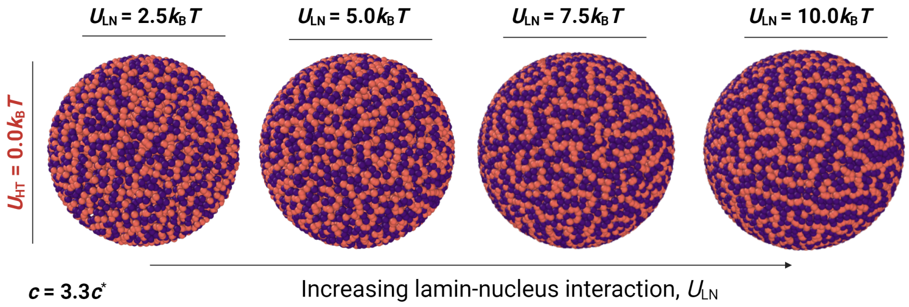


FIG. S15: Organized domain formation can be observed with $U_{\mathrm{LN}}$ alone at high concentrations. The effect of lamin-nucleus association potential, $U_{\mathrm{LN}}$, in the absence of lamin-lamin association potential, $U_{\mathrm{HT}}$, at fixed concentration, $c = 3.3c^*$.

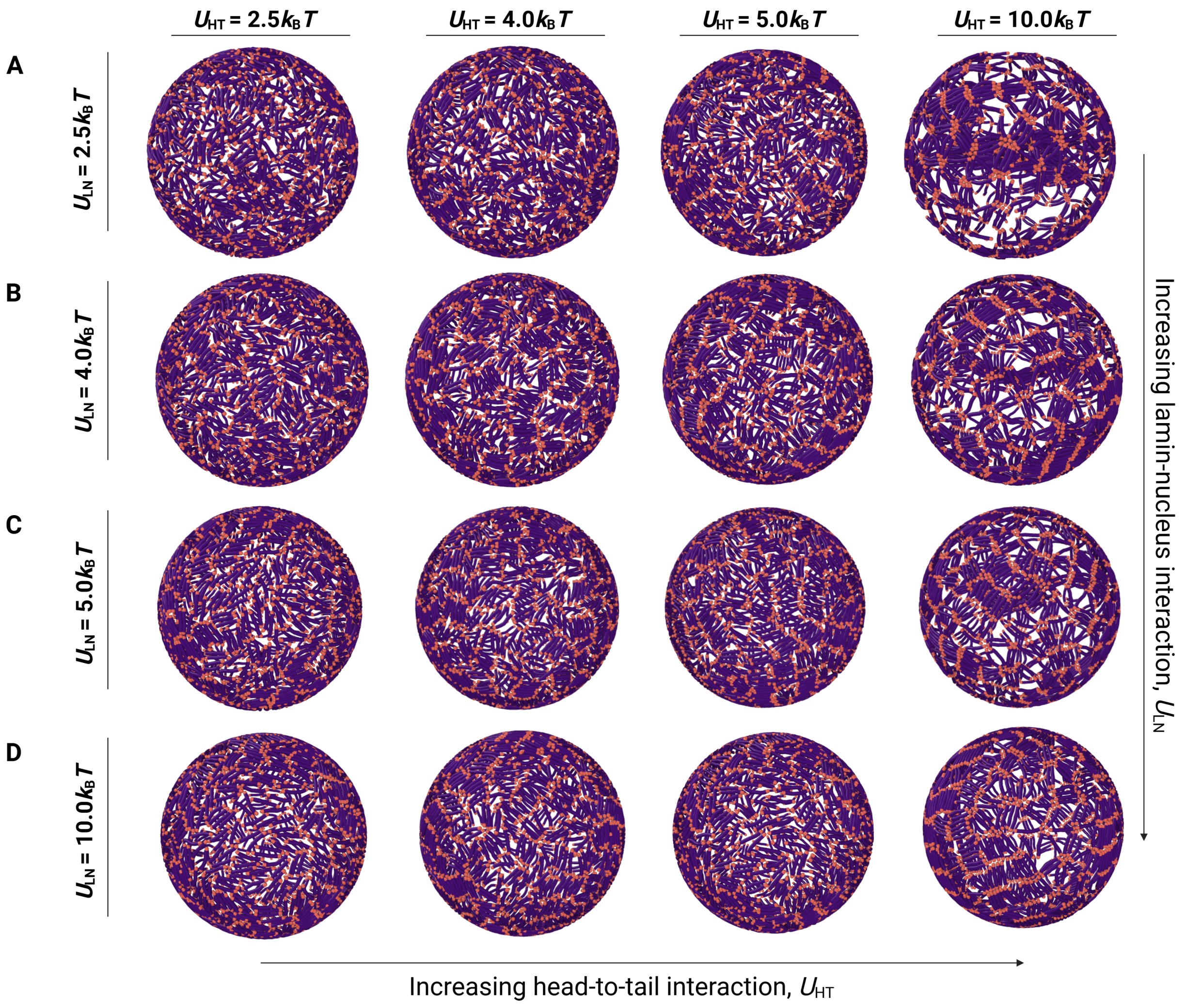


FIG. S16: Lamin fibers exhibit a more isotropic and/or network-like configuration instead of nematic domains when $c \leq c^*$. Snapshots for $c \leq 2c^*$ case i.e. $c = 1.6c^*$ showing the effect of $U_{\mathrm{HT}}$ for $U_{\mathrm{LN}}$ = **A)** 2.5, **B)** 4.0, **C)** 5.0, and **D)** 10.0$k_{\mathrm{B}}T$.

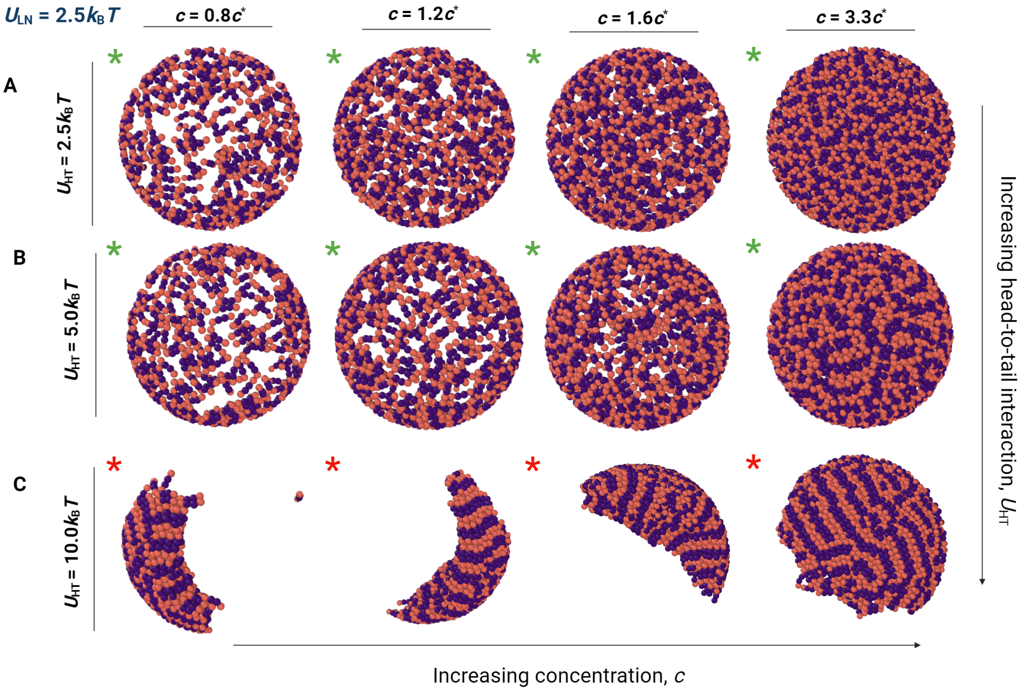


FIG. S17: Exterior cross-sections of the nuclear lamina at $U_{\mathrm{LN}} = 2.5k_{\mathrm{B}}T$ for 3 different lamin-lamin attraction strengths, $U_{\mathrm{HT}}$ at various lamin concentrations for $n = 4$ bead lamins.

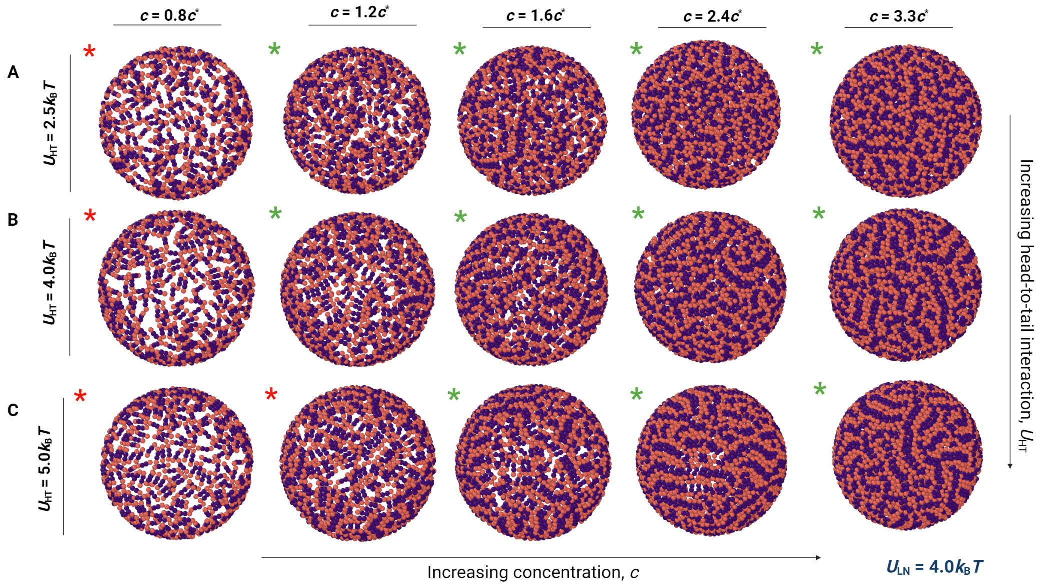


FIG. S18: Exterior cross-sections of the nuclear lamina at $U_{\mathrm{LN}} = 4.0k_{\mathrm{B}}T$ for 3 different lamin-lamin attraction strengths, $U_{\mathrm{HT}}$ at various lamin concentrations for $n = 4$ bead lamins.

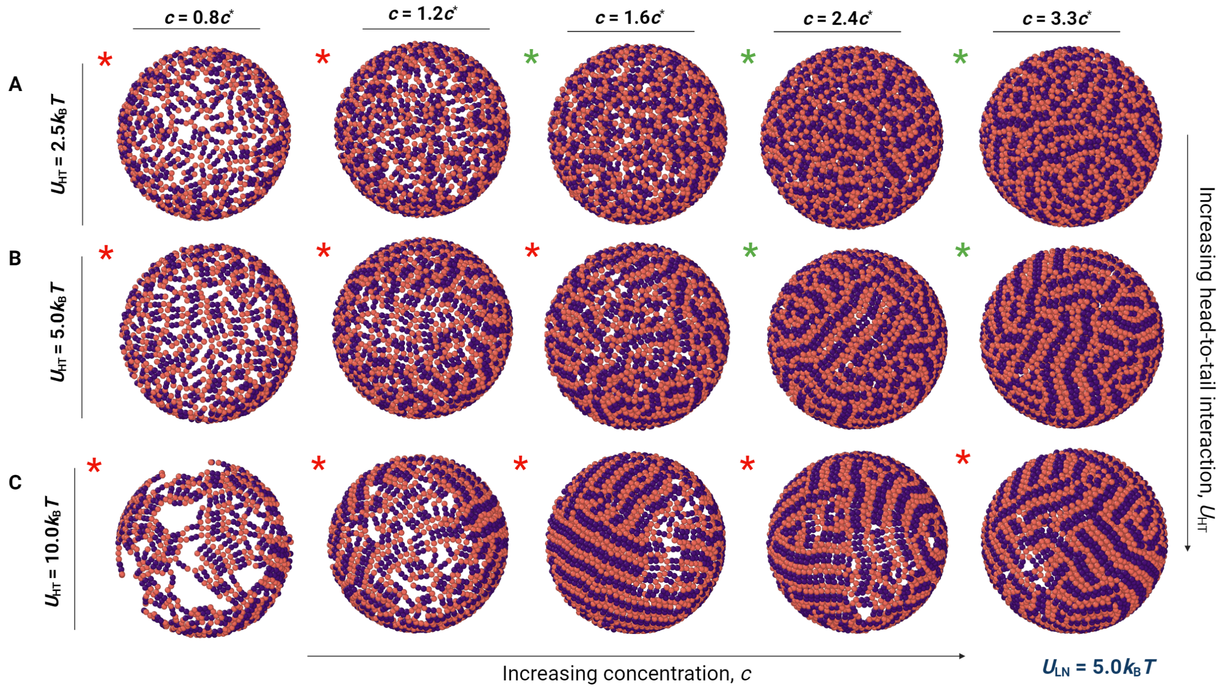


FIG. S19: Simulation snapshots of the nuclear lamina exterior at $U_{\mathrm{LN}} = 5.0 k_{\mathrm{B}} T$ for 3 different lamin-lamin attraction strengths, $U_{\mathrm{HT}}$ at various lamin concentrations for $n = 4$ bead lamins.

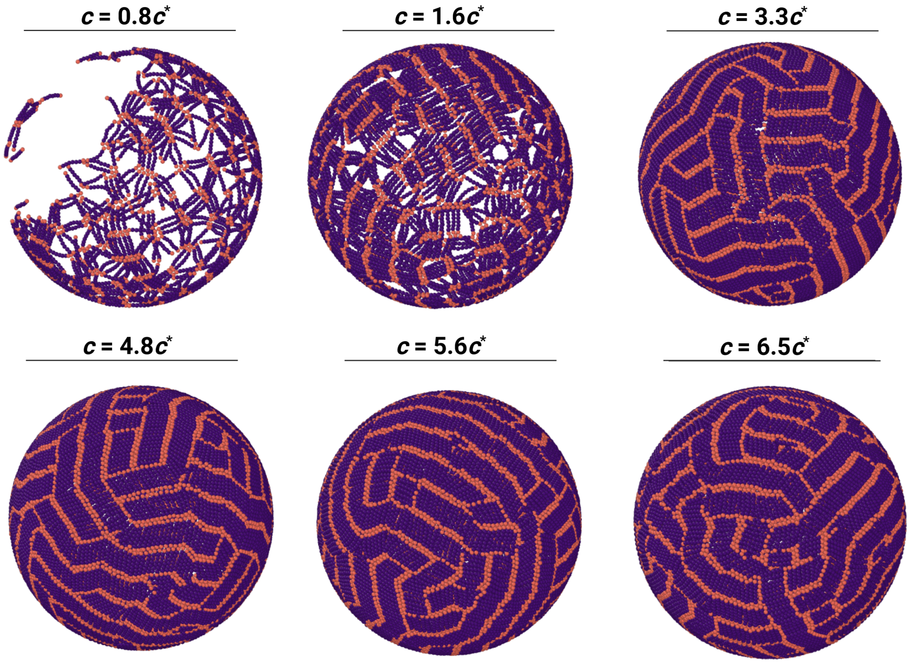


FIG. S20: $U_{\mathrm{LN}} = 5.0 k_{\mathrm{B}} T$ is strong enough to localize 8-bead lamin fibers in a nematic fashion at the surface when $c >> c^{*}$.

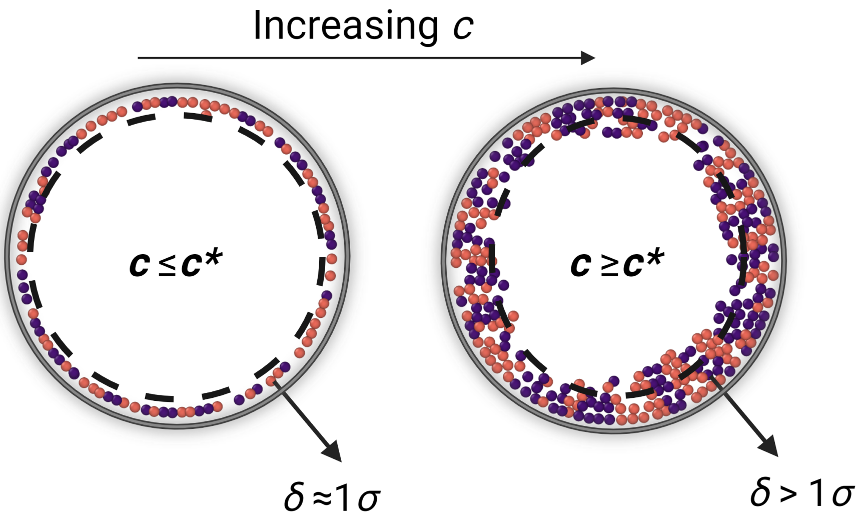


FIG. S21: The effect of concentration on the thickness of the nuclear lamina when $U_{\rm HT} = U_{\rm LN} = 10.0 k_{\rm B}T$. Thickness, $\delta$, exceeds a single monolayer when $c \geq c^*$.

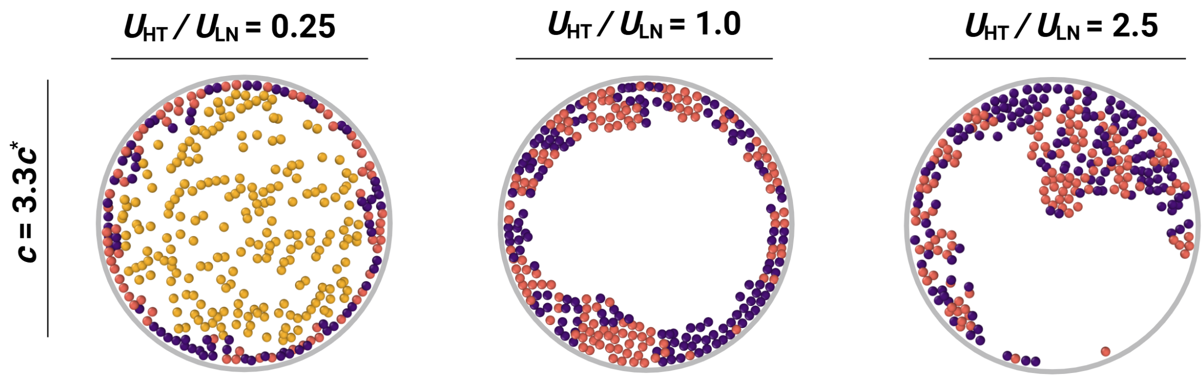


FIG. S22: $1\sigma$ wide slices showing various thickening configurations that can be obtained, dictated by the lamin-lamin interaction strength, $U_{\rm HT}$. $U_{\rm LN} = 10.0 k_{\rm B}T$ and concentration is fixed to $c = 3.3c^*$ for $n = 4$ bead lamin fibers. Trends for $n = 8$ bead lamin fibers are shown in the main text.

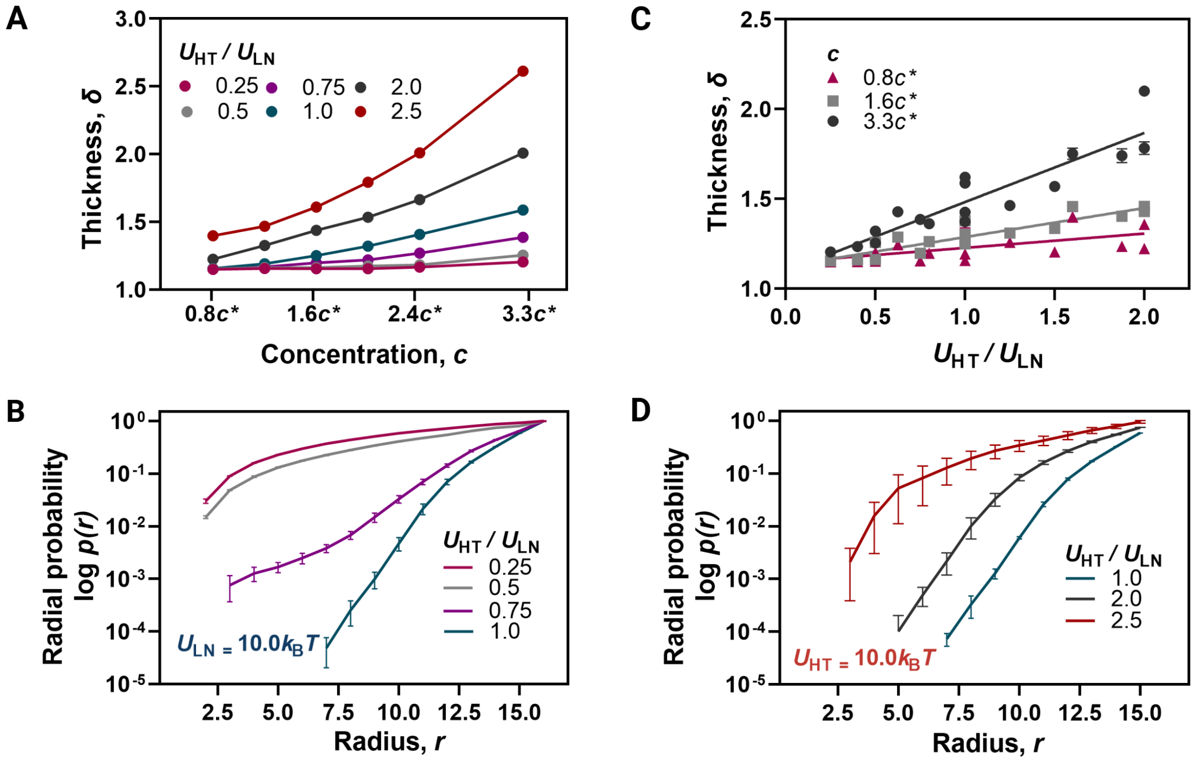


FIG. S23: Trends observed for lamina thickness for $n = 4$ bead lamin fibers. **A)** Nuclear lamina thickness, $\delta$, vs. concentration, $c$, at various $U_{\mathrm{HT}}/U_{\mathrm{LN}}$. **B** Logarithm of radial probability distribution, *p(r)*, as a function of radial distance, $r$, at 4 different $U_{\mathrm{HT}}/U_{\mathrm{LN}}$ for $c = 3.3c^*$ and $U_{\mathrm{LN}} = 10.0k_{\mathrm{B}}T$. **C)** Thickness, $\delta$, vs. ratio of affinities, $U_{\mathrm{HT}}/U_{\mathrm{LN}}$, at 3 concentrations i.e., $c = 0.8c^*$, $1.6c^*$, and $3.3c^*$ with over 20 data points. **D)** Logarithm of radial probability distribution, *p(r)*, as a function of radial distance, $r$, at 3 different $U_{\mathrm{HT}}/U_{\mathrm{LN}}$ for $c = 3.3c^*$ and $U_{\mathrm{HT}} = 10.0k_{\mathrm{B}}T$.

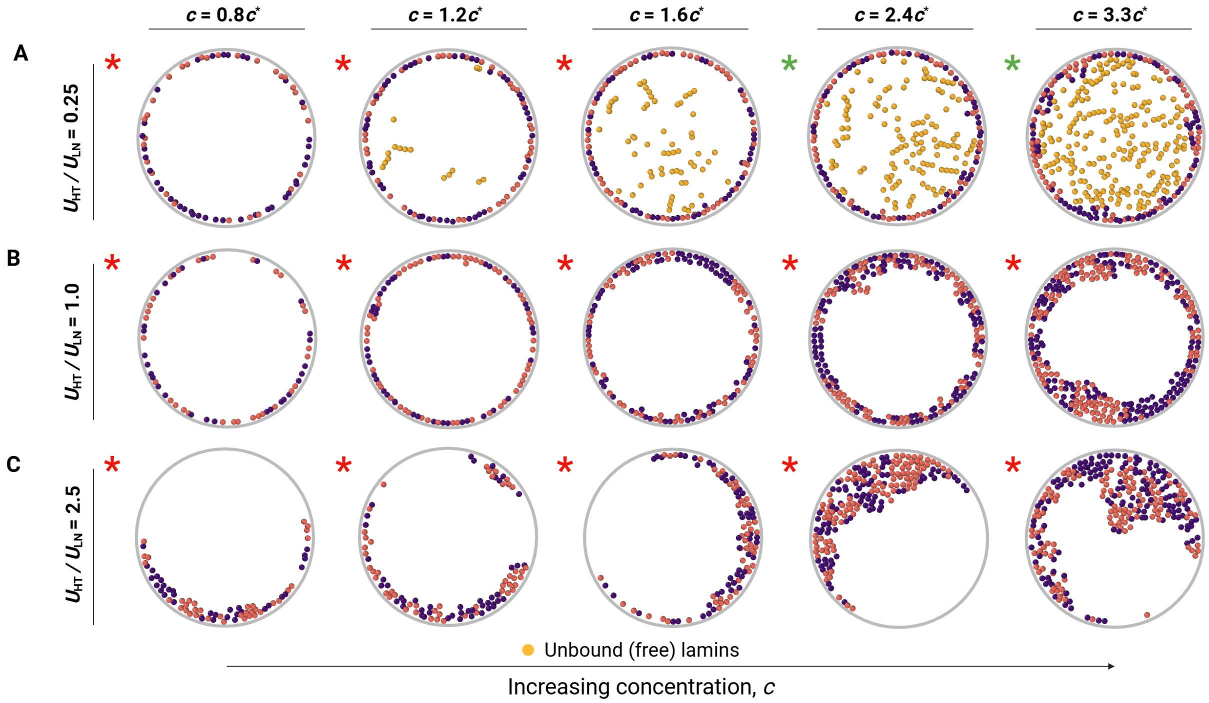


FIG. S24: Snapshots of $1\sigma$ wide slices from the center of the model nucleus at various lamin concentrations for $U_{\rm HT}/U_{\rm LN}$ = **A)** 0.25, **B)** 1.0, and **C)** 2.5. Yellow chains represent *unbound* lamin proteins participating in the kinetic exchange.

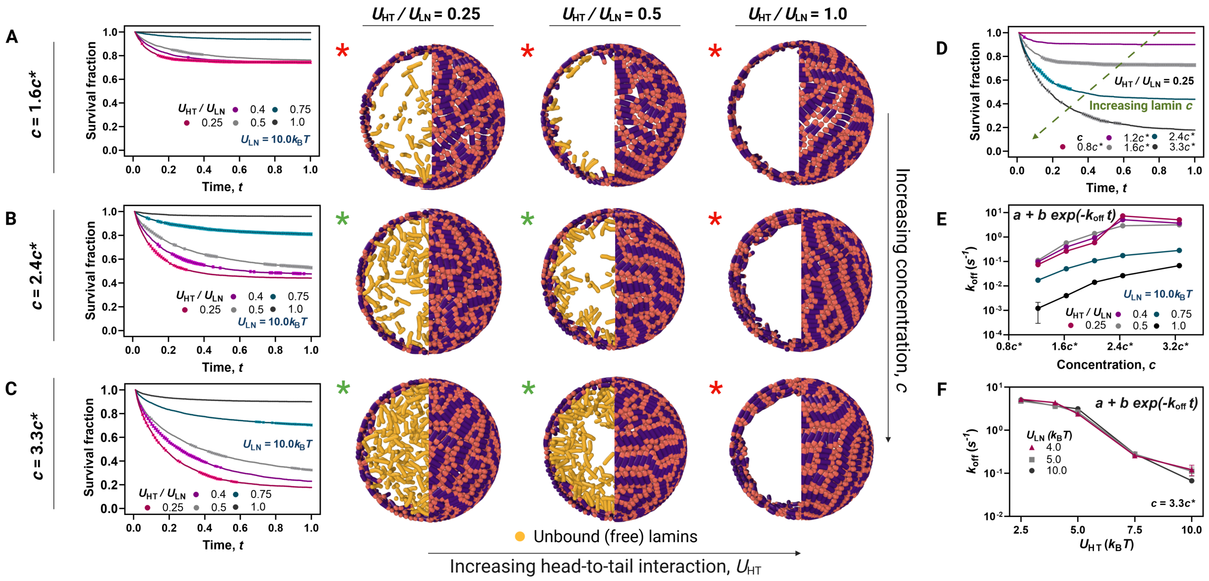


FIG. S25: The dependence of lamin dissociation kinetics on concentration, $c$, and ratio of head-to-tail association potential to lamin-shell affinity, $U_{\mathrm{HT}}/U_{\mathrm{LN}}$ for 4-bead lamins. Graphs on the left represent the behavior of the normalized survival fraction, $n(t)/n_0$, of *bound* nuclear lamins over time, $t$. The solid curves are the exponential fits (Eq. **??**). Snapshots in the middle show $10\sigma$ wide slices from the center of the model nucleus. Yellow chains represent *unbound*, free lamin proteins participating in a kinetic exchange. **A-C)** Illustrations representing the degree of exchange between nucleoplasmic and shell-bound lamins at three different concentrations: $c =$ **A)** $1.6c^*$, **B)** $2.4c^*$, and **C)** $3.3c^*$ at fixed $U_{\mathrm{LN}} = 10.0k_{\mathrm{B}}T$. **D)** Time-dependent decay curve for lamin dissociation at various concentrations to extract off-rates, $k_{\mathrm{off}}$, from the fraction of remaining *bound* lamins over time with fixed $U_{\mathrm{HT}}/U_{\mathrm{LN}} = 0.25$. **E-F)** Off-rates extracted from the survival fraction data. **E)** Off-rates as a function of lamin concentration for various $U_{\mathrm{HT}}/U_{\mathrm{LN}}$ at fixed $U_{\mathrm{LN}} = 10.0k_{\mathrm{B}}T$. **F)** Off-rates as a function of head-tail association potential, $U_{\mathrm{HT}}$, for various $U_{\mathrm{LN}}$ at a fixed concentration, $c = 3.3c^*$.

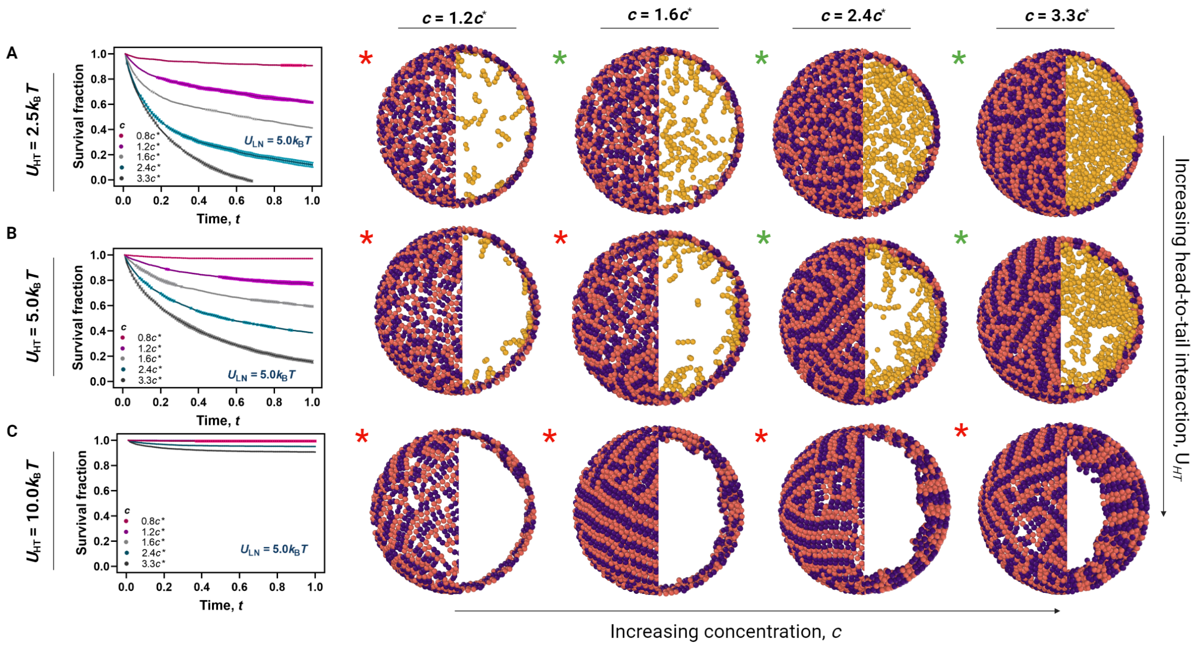


FIG. S26: Lamin dissociation curves, and simulation exterior/ interior simulation snapshots at various concentrations and head-to-tail association potentials, $U_{\mathrm{HT}}$, when $U_{\mathrm{LN}}$ $= \mathbf{5.0}k_{\mathrm{B}}T$. Yellow chains represent exchanging lamins. Red asterisks (*) represent cases where lamins are restricted at the nuclear periphery, while green asterisks (*) represent the dynamic exchange regime. **A)** Dynamic lamin exchange is observed when $c \geq 1.5c^{*}$ when lamin-lamin association potential is low i.e. $U_{\mathrm{HT}} = 2.5k_{\mathrm{B}}T$. **B)** Lamin exchange is restricted for $c = 1.6c^{*}$ when $U_{\mathrm{HT}} = 5.0k_{\mathrm{B}}T$. **C)** No lamin dissociation and nuclear lamina thickening is observed when $U_{\mathrm{HT}} = 10.0k_{\mathrm{B}}T$. In summary, similar trends are observed in the interior as $U_{\mathrm{LN}} = 10.0k_{\mathrm{B}}T$, shown in the main text (Fig. 2).

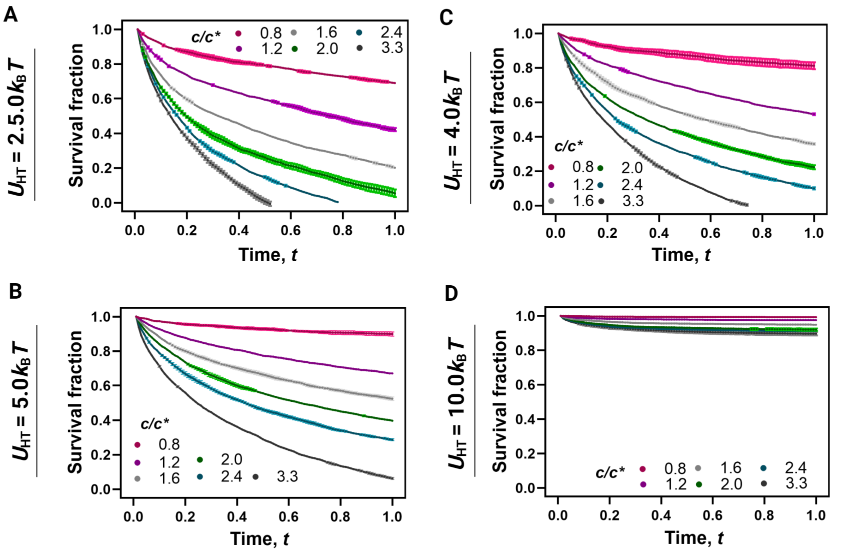


FIG. S27: Comparison of the survival fractions of lamin proteins at $U_{\rm HT}$ = **A)** 2.5, **B)** 4.0, **C)** 5.0, and **D)** 10.0$k_{\rm B}T$. Lamin-nucleus association potential is fixed at $U_{\rm LN}$ = $\mathbf{4.0}k_{\rm B}T$. While a similar general dissociation trend is observed at higher concentrations, the survival fractions decay faster at all concentrations than when $U_{\rm LN} = 5.0$, and $10.0k_{\rm B}T$.

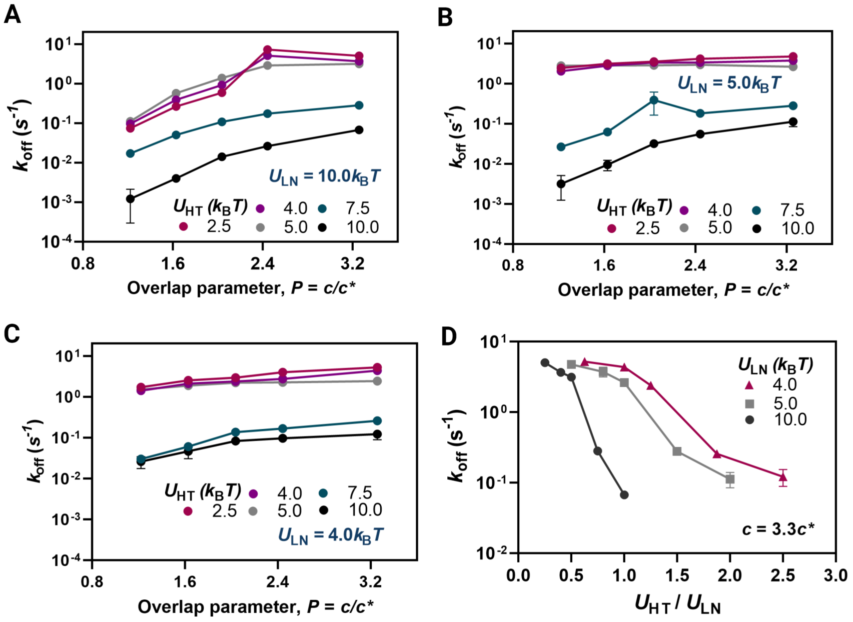


FIG. S28: Effect of $U_{\rm LN}$, $U_{\rm HT}$, and $c$ on lamin dissociation off-rates, $k_{\rm off}$. Off-rates with respect to overlap parameter, $P$, at $U_{\rm LN}$ = **A)** 10.0, **B)** 5.0, and **C)** 4.0$k_{\rm B}T$. **D)** Off-rates with respect to various affinity ratios, $U_{\rm HT}/U_{\rm LN}$ at fixed concentration, $c = 3.3c^*$.

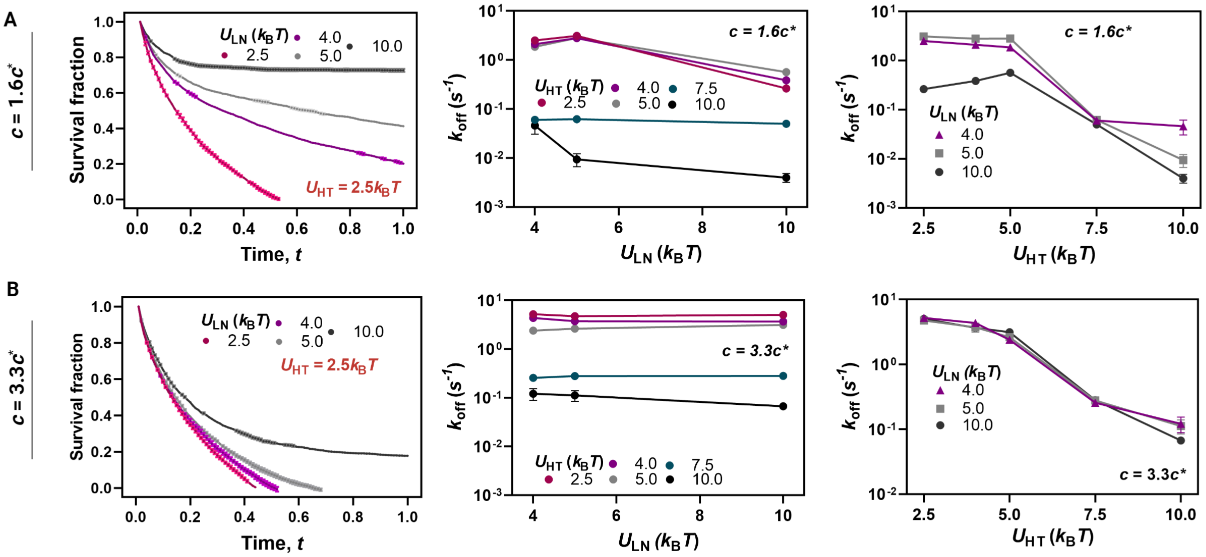

FIG. S29: Head-to-tail interaction potential, $U_{\mathrm{HT}}$, seems to have a more significant effect on lamin dissociation kinetics than the association of the lamin fibers to the nuclear periphery, $U_{\mathrm{LN}}$, particularly when $c > 2c^*$. Comparison of survival fraction, and off-rates vs. $U_{\mathrm{LN}}$, and $U_{\mathrm{HT}}$ at intermediate, **A)** $c = 1.6c^*$, and high, **B)** $c = 3.3c^*$, concentrations.

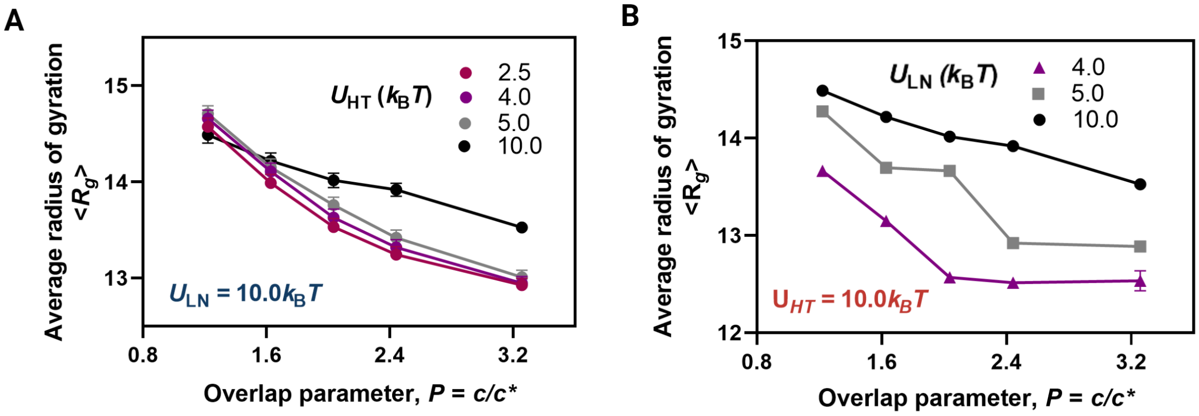

FIG. S30: The effect of $U_{\mathrm{HT}}$, $U_{\mathrm{LN}}$, and concentration, $c$, on the average radius of gyration of lamin proteins, $\langle R_g \rangle$. **A)** $\langle R_g \rangle$ vs. Overlap parameter, $P$, at 4 different head-to-tail association potentials. **B)** $\langle R_g \rangle$ vs. Overlap parameter, $P$, at 3 lamin-nucleus association potentials. $\langle R_g \rangle$ decreases as $U_{\mathrm{LN}}$ decreases, indicating that lamin fibers are more spread out towards the interior at lower lamin-nucleus association potentials.
